# Bottom-Illuminated Orbital Shaker for Microalgae Cultivation

**DOI:** 10.1101/2020.05.01.071878

**Authors:** Jakub Nedbal, Lu Gao, Klaus Suhling

## Abstract

A bottom-illuminated orbital shaker designed for the cultivation of microalgae suspensions is described in this open-source hardware report. The instrument agitates and illuminates microalgae suspensions grown inside flasks. It was optimized for low production cost, simplicity, low power consumption, design flexibility, consistent, and controllable growth light intensity.

The illuminated orbital shaker is especially well suited for low-resource research laboratories and education. It is an alternative to commercial instruments for microalgae cultivation. It improves on typical do-it-yourself microalgae growth systems by offering consistent and well characterized illumination light intensity. The illuminated growth area is 20 cm × 15 cm, which is suitable for three T75 tissue culture flasks or six 100 ml Erlenmeyer flasks. The photosynthetic photon flux density, is variable in eight steps (26 – 800 μmol · m^−2^ · s^−1^) and programmable in a 24-hour light/dark cycle. The agitation speed is variable (0 – 210 RPM). The overall material cost is around £300, including an entry-level orbital shaker. The build takes two days, requiring electronics and mechanical assembly capabilities. The instrument build is documented in a set of open-source protocols, design files, and source code. The design can be readily modified, scaled, and adapted for other orbital shakers and specific experimental requirements.

The instrument function was validated by growing fresh-water microalgae *Desmodesmus quadricauda* and *Chlorella vulgaris*. The cultivation protocols, microalgae growth curves, and doubling times are included in this report.

**Specifications table:** 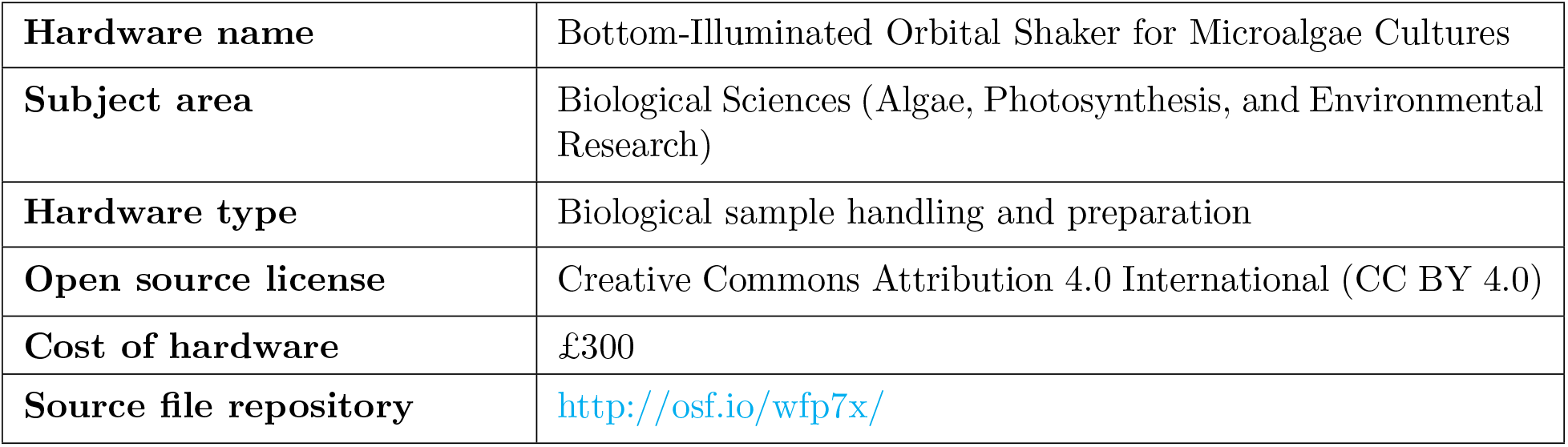

## 1. Hardware in context

Microalgae, like plants, use photosynthesis as the primary energy source for their metabolic needs [1]. In laboratory conditions, microalgae are grown using dedicated instruments in growth media, which provide the nutrients and water [2, 3]. These instruments vary in size between microfluidic [4, 5] and industrial scale implementations [3, 6]. Their purpose is to agitate and illuminate the cultures. Agitation is done by shaking, stirring or gas sparging to ensure microalgae mixing, nutrient, and gas exchange. The illumination typically covers the photosynthetically active spectral range 400 nm − 700 nm (white), which excites a range of endogenous fluorophores required for full metabolic activity. Their illumination intensity is variable and programmable for daily (diurnal) light/dark cycle.

The instrument described in this report is an orbital shaker with a bottom-mounted light source for growing 10s − 100s ml of microalgae suspensions in flasks (Figure 1A). The instrument offers consistent and controllable illumination with low power consumption at a fraction of the cost of a commercial instrument. Laboratory microalgae cultivation is typically done in photobioreactors, illuminated cabinets or on shakers. Closed-system photobioreactors (Figure 1B) offer controlled cultivation conditions for high biomass production [7, 8]. Illuminated cabinets with controlled environment (temperature, humidity, gas composition) are used to grow microalgae inside flasks on orbital shakers (Figure 1C). In the simplest case, this specialized cabinet is replaced by a common top-mounted lamp illuminating the microalgae culture grown at room temperature on an orbital shaker (Figure 1D).

**Figure 1:**
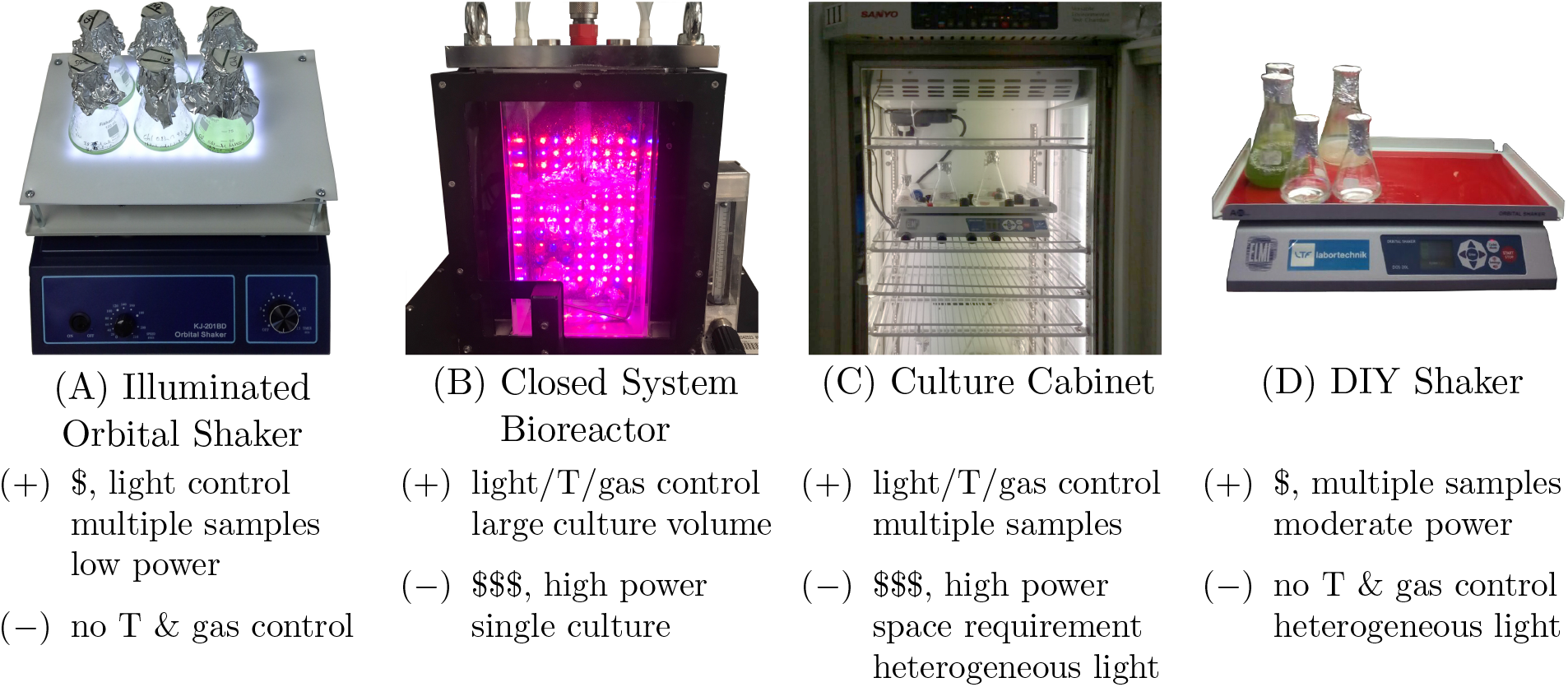
Different cultivation systems for microalgae and a summary of their strengths and weaknesses. A) Illuminated orbital shaker described in this manuscript. B) Closed system bioreactor for high biomass volume; C) Culture cabinet for large number of different cultures; D) Low-cost DIY shaker under top-mounted lights. T stands for temperature, $ for low cost, and $$$ for high cost.

Closed system bioreactors (Figure 1B) and temperature-controlled illuminated growth cabinets (Figure 1C) are designed to offer regulated and versatile conditions for microalgae cultivation [2, 3]. However, these specialized machines are expensive (£1000s − £10000s), can have large power (> kW) and space requirements, and operating costs. Yet, these commercial systems are not always essential and do-it-yourself (DIY) solutions to microalgae cultivation are therefore common [9, 10, 11, 12, 8]. However, the technical and design details of DIY cultivation systems are seldom published and open source hardware (OSHW) publications on microalgae cultivation remain rare. An OSHW photobioreactor design, build instructions, and application protocols were published recently in a peer-reviewed article [8]. ‘NinjaPBR’ is another photobioreactor for microalgae cultivation with its design files, assembly instructions, and control software made available on Github [13]. ‘Bioreactor’ is a device for the cultivation of bacteria, currently under development, released on GitHub [14]. It lacks a light source required for phototrophic cultivation of microalgae, but it lends itself to this modification. A suitable light source with programmable power, spectrum, and period is described in [15]. Another exemplary OSHW device is an automated turbidostat, which monitors and maintains optimal microalgae culture density to maximize photobioreactor production yield [16, 17].

According to Büchs, over 90 % of all culture experiments in biotechnology are performed in shaking bioreactors [18]. Similarly, in laboratory cultivation of microalgae, orbital shakers are widely used. Typically, flasks with microalgae are placed on top of an orbital shaker (Figure 1D) with a light source illuminating the culture from the top [9, 10]. This solution is cheap and simple to implement. However, it cannot ensure consistent and predictable illumination of the microalgae suspensions inside the flasks. The flask lids may screen off the light and cast shadows and the lamp illumination may not be homogeneous across the area of the shaker. Bottom-illuminated orbital shaker systems for microalgae cultivation offer more consistent illumination [11, 12], but are not widely used. This manuscript describes an open hardware design for a bottom-illuminated orbital shaker for the cultivation of microalgae (Figure 1A). It was optimized for production cost simplicity, low power consumption, design flexibility, and consistent and controllable growth light intensity to create reproducible experimental conditions.

## 2. Hardware description

This work describes an illuminated orbital shaker built around a commercial orbital shaker. A custom light-emitting diode (LED) [19, 20] array illuminator and an electronic LED controller are placed on top of the shaking platform. An elevated transparent platform made of clear acrylic is fixed over the LED illuminator. The microalgae cultures are placed on top of this elevated platform, being illuminated from the bottom by the LEDs, and agitated by the rotational motion of the orbital shaker.

The LED illuminator is a light source with a rectangular area of 20 cm × 15 cm, positioned only 15 mm below the microalgae culture. The close proximity of the light source and its positioning below the culture flasks is highly beneficial. The illumination of the culture is consistent over time, regardless of the type of culture flask used, and requires comparably low power consumption (≤ 29 W) to achieve irradiance sufficiently high for majority of microalgae cultivation requirements. The light/dark cycle is set by programming a 24-hour socket timer, which regularly turns the light on and off. The irradiance at the bottom interface of the microalgae culture is reproducibly adjustable in eight steps, using a custom electronics controller described in Section 5. The brightness of the light can be regulated by manual switches or an external microcontroller. Excess heat generated by the LEDs is removed by a fan-cooled aluminium heatsink. The cooling ensures that the temperature at the top of the illuminated shaker platform does not increase more than 1 °C above the ambient temperature even at the highest light output.

The bottom-illuminated orbital shaker function was validated by successful cultivation of freshwater microalgae, described in Section 7.2. However, its design is more versatile and should support wider range of applications than described in this report:

- Different species of freshwater and seawater suspension microalgae and cyanobacteria can be cultivated in the appropriate growth medium [9, 21, 10].
- Thermophilic or psychrotrophic microalgae and cyanobacteria can be cultivated with the orbital shaker placed inside a temperature-controlled space [22, 23, 24].
- Seed cultures can be cultivated for inoculation into high-volume bioreactors [9].
- The design can be modified for different orbital shaker models and types. A readily available shaker can be substituted for the model described here. The design can be modified to feature larger illuminator and orbital shaker to support larger culture volumes.
- The peak brightness of the illuminator can be changed by altering the spacing of the LED strips^1^.
- The spectral properties of the light source can be modified to suit different experimental regimes [25]. The white LED strips, described here, can be swapped for multi-color LED strips[15]. Multiple copies of the described LED controller can be built to independently control the output of the three colors of the multicolor LED strip, and thus vary the spectral properties and timing of the light source.
- The illumination light intensity and cycle can be controlled using a microcontroller or a computer for advanced growing protocols. The described LED controller is prepared for such external control. It has 0.30 V − 1.25 V analog input regulating the LED current between 25 % − 100 %. Arduino DUE, Zero or the MKR-family feature in-built digital-to-analog converters for direct voltage control of LED illuminator brightness.
- Pulse width modulation (PWM) can be used to control the brightness with most microcontrollers, including all members of the Arduino and Raspberry Pi families using the same external input of the LED controller.
- The light/dark cycle can be upgraded to longer and more complex illumination patterns by a straightforward replacement of the basic 24-hour plug-in timer with a more advanced weekly programmable plug-in timer.
- The orbital shaker can be used in education. It could be an educational engineering project focusing on the instrument development, and a tool for learning about population biology and photosynthesis [26].

## 3. Design files

Design files are provided for readers to use them directly or modify them according to their specific needs. The design files and the associated assembly steps can be grouped into four areas:

- LED controller electronics.
- 3D printed case for the LED controller electronics.
- Cooled LED illuminator.
- Transparent orbital shaker platform.

The electronic circuit and the printed circuit board (PCB) have been designed in KiCAD electronic design automation suite. The 3D and 2D computer aided design (CAD) models of the electronics circuit case and the transparent orbital shaker platform were designed in Onshape CAD software system.

### 3.1 Design files summary

The design files are listed in Table 1 and are briefly described in the list below. Their use in the build of the illuminated orbital shaker in explained later in Section 5 and the accompanying protocols.

- `LEDcontroller.zip` The archive contains KiCAD project source files with the electronics schematics, printed circuit board (PCB), bill-of-materials list, and PCB production files. The latest version is available at rebrand.ly/etuuxu. The schematics of the LED controller is in Supplementary Figure S1.
- `LEDcontroller_PCB.zip` Gerber files for LED controller PCB production are in this archive. The latest version is available at rebrand.ly/xhsc9i.
- `LEDcontroller_Case.zip` STL files for 3D printing the parts of the custom LED controller case are in this archive. The Onshape 3D CAD project with the 3D models of the electronics and the case is available at rebrand.ly/hvjd1o.
- `Acrylic_Sheet.dxf` This DXF file is for the use with laser cutters to produce the secondary transparent orbital shaker platform from clear acrylic. The Onshape 3D CAD project with the latest DXF file, technical drawing and 3D model of the shaker platform is available at rebrand.ly/9nqpar.

**Table 1:** List of design files used in the build of the illuminated orbital shaker for microalgae cultivation.

| Design filename | File type | Open source license | Location of the file |
| --- | --- | --- | --- |
| LEDcontroller.zip | KiCAD project | CC BY-SA 4.0 | osf.io/b4wph |
| LEDcontroller_PCB.zip | Gerber PCB files | CC BY-SA 4.0 | osf.io/vejbw |
| LEDcontroller_Case.zip | 3D CAD files | CC BY-SA 4.0 | osf.io/dsmk7 |
| Acrylic_Sheet.dxf | Laser cutter file | CC BY-SA 4.0 | osf.io/qwh6t |

## 4. Bill of materials

The materials, required to build the bottom-illuminated orbital shaker for microalgae cultivation, include an orbital shaker, mechanical fasteners and fixings, clear acrylic sheet, electrical, and electronics components. A number of workshop tools and stationaries are used in the process of building the illuminated orbital shaker. Both the components and tools are organized in a spreadsheet at osf.io/tqhy9 and are also listed in Supplementary Tables S3 to S7.

The spreadsheet is divided into five tabs named:

- **Electronics Parts:** Electronics components are soldered to the PCB to complete the LED controller circuit. Their total cost is £38.
- **Laboratory Parts:** The orbital shaker forms the basis of this project. A commonly available low cost orbital shaker (£95) is described in this manuscript. However, other makes and models can be substituted - including used ones.
- **Fixings:** Widely available screws, nuts, washers, and standoffs are required. The total cost is £9.
- **LED Illuminator Parts:** Electrical and electromechanical components required to build the LED illuminator include self-adhesive LED strips, heatsink, cooling fans, cables, power supply, 24-hour socket time switch, and a clear acrylic sheet. The total cost is £155.
- **Tools:** A number of commonly available workshop tools and office stationaries are required during the assembly of the orbital shaker. The total cost of brand new tools used in the assembly would be £436, excluding the 3D printer. The tools are organized into five categories:

– **Workshop Tools:** Common mechanical workshop tools, thread tap set, tap wrench, and a digital multimeter.
– **Stationaries:** Multipurpose glue, scissors, fine-tip marker pen, and a ruler.
– **Soldering equipment:** Soldering station with solder and flux, electrical tape, and PCB-cleaning supplies (isopropyl alcohol, tub, and brush).
– **3D Printing:** Fused deposition modeling 3D printer with consumables or a commercial 3D printing service.
– **Cutting and Drilling Tools:** Either a drill, drill bit, hacksaw, and a sandpaper, or a laser cutter capable of cutting acrylic sheets.

The electronics circuit PCB manufacture can be outsourced. The total cost of the manufacture, including shipping, is £10. The instructions for ordering the custom PCB are in the following Section 5.

The total cost of the bill of materials, including surplus electronics parts and fixings, is approximately £300. This price includes a basic shaker for ≈ £100.

## 5. Build instructions

Building the bottom-illuminated orbital shaker for microalgae cultivation requires electronics and mechanical assembly skills. The build is thoroughly documented in a series of protocols listed below. The protocols are published online using **protocols.io** platform under CC BY 4.0 license. The links to the protocols and their brief summaries are in Table 2. Their more detailed description follows.

- **Protocol 1: Illuminated Orbital Shaker for Microalgae Culture (osf.io/hd2c6)** [27] This is a high-level protocol summarizing the steps in building the illuminated orbital shaker. It starts with procuring parts and tools and finishes with the the final assembly. Links to the sub-protocols, explaining each step in detail, are provided.
- **Protocol 2: Procuring Parts for Algal Shaker (osf.io/jy7gc)** [28] Number of tools, equipment and parts are required to build the illuminated orbital shaker. Detailed bills of electronics and mechanical materials are provided. The process of ordering the printed circuit board for the LED controller is also detailed here.
- **Protocol 3: Assembling LED Controller Electronics (osf.io/bftxm)** [29] The LED controller is an electronics circuit, which regulates the illuminator power by varying LED current and drives the cooling fans, which prevent overheating of the LEDs and microalgae cultures. The function of the LED controller electronics circuit, step-by-step assembly, and testing instruction are detailed in this protocol. A 3D render of the assembled electronics circuit is shown in Figure 2A.
- **Protocol 4: 3D Printing Case for LED Controller (osf.io/7ycnr)** [30] The design files to produce a custom housing for the LED controller electronics by 3D printing are introduced. The process of the assembly of the LED controller is explained in this protocol. A 3D render of the assembled LED controller inside the case is shown in Figure 2B.
- **Protocol 5: Assembling Cooled LED Illuminator (osf.io/hywec)** [31] The cooled LED illuminator consists of a heatsink holding the illuminating LED strips, cooling fans, and the LED controller. Instructions for the mechanical and electrical assembly of the cooled LED illuminator are provided in this protocol. The initial testing of the LED controller circuit is also outlined here. A photograph of the assembled LED illuminator is shown in Figure 2D.
- **Protocol 6: Cutting and Drilling Clear Acrylic Sheet (osf.io/69f8t)** [32] A clear acrylic sheet is used as a secondary transparent shaking platform holding the microalgae culture flasks. Two sets of instructions are provided in this protocol: One for the manual cutting of the clear acrylic sheet using workshop tools; the second for automatic cutting using a laser cutter. A 3D render of the cut and drilled acrylic sheet is in Figure 2C.
- **Protocol 7: Assembling Algal Shaker (osf.io/ewc87)** [33] Once all components of the illuminated orbital shaker are built, the final instrument is assembled by mounting the parts to the orbital shaker platform and connecting them electrically. These last assembly steps are detailed in this protocol. A photograph of the assembled illuminated orbital shaker is shown in Figure 2E.
- **Protocol 8: Measuring PPFD on Algal Shaker (osf.io/va54p)** [34] The light output is characterized by measuring the photosynthetic photon flux density (PPFD), as described in this protocol. Experimental details are in Section 7.1.

**Figure 2:**
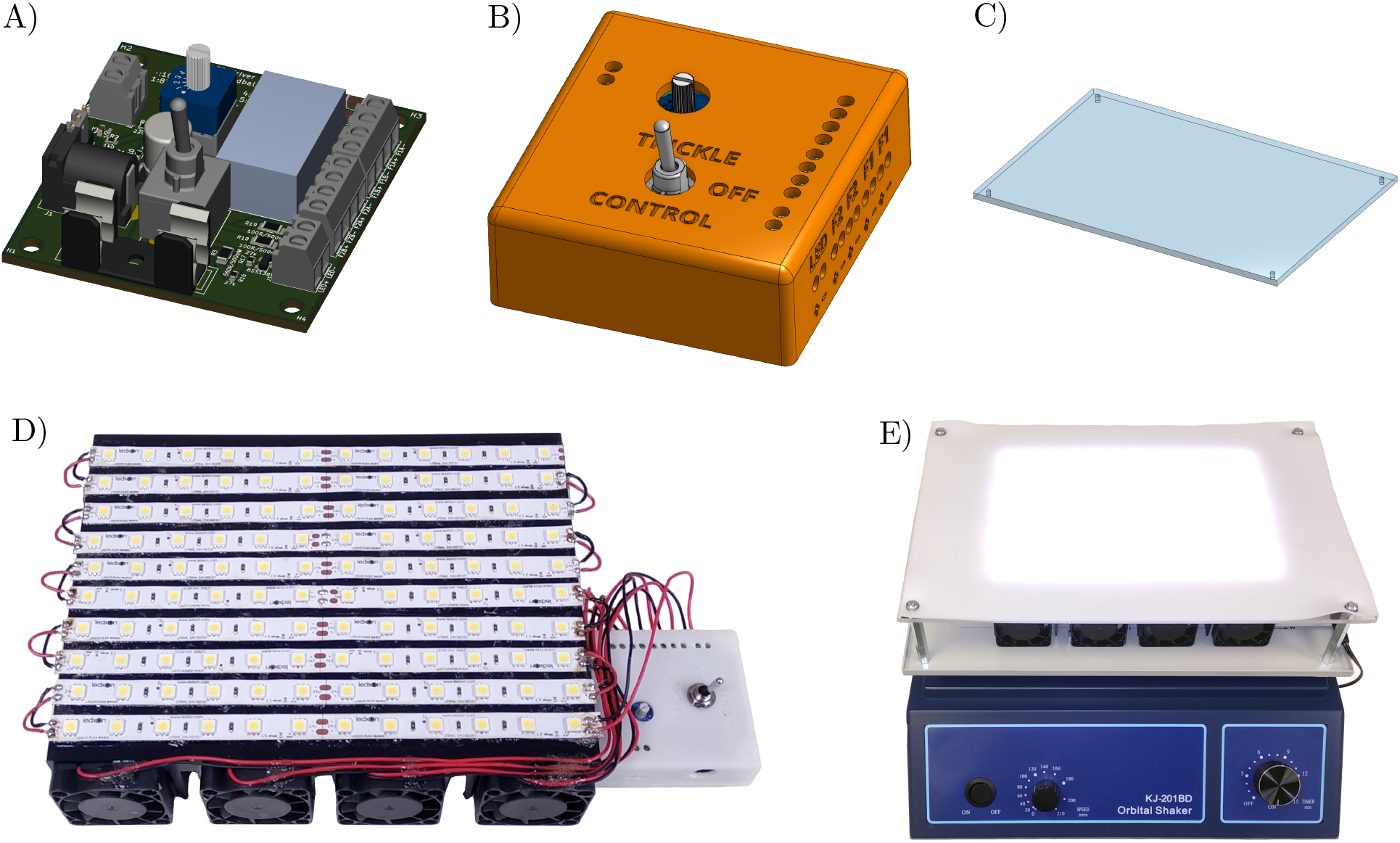
Progress of building the illuminated orbital shaker. A) LED controller electronics circuit; B) 3D printed case for LED controller; C) Clear acrylic sheet for transparent shaker platform; D) LED illuminator; E) Assembled illuminated orbital shaker.

**Table 2:**
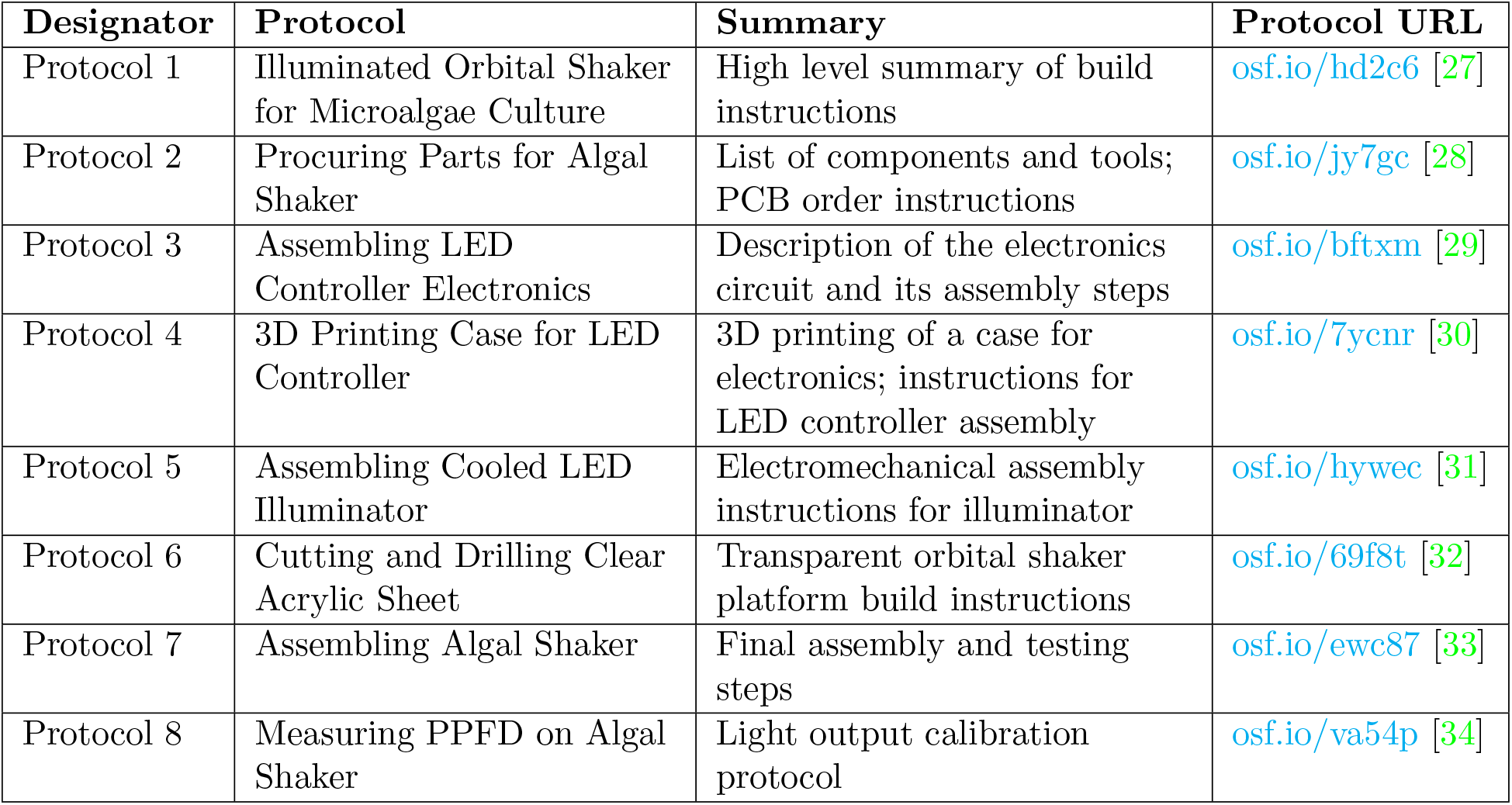
List of protocols with instructions on building and calibrating the illuminated orbital shaker platform. All protocols are licensed under CC BY 4.0 license. The URL links to the protocols in this table lead to their static snapshots current at the time of this publication. The bibliographic citations point to entries on **protocols.io**, which may be updated with new versions in the future.

| Designator | Protocol | Summary | Protocol URL |
| --- | --- | --- | --- |
| Protocol 1 | Illuminated Orbital Shaker for Microalgae Culture | High level summary of build instructions | osf.io/hd2c6 [27] |
| Protocol 2 | Procuring Parts for Algal Shaker | List of components and tools; PCB order instructions | osf.io/jy7gc [28] |
| Protocol 3 | Assembling LED Controller Electronics | Description of the electronics circuit and its assembly steps | osf.io/bftxm [29] |
| Protocol 4 | 3D Printing Case for LED Controller | 3D printing of a case for electronics; instructions for LED controller assembly | osf.io/7ycnr [30] |
| Protocol 5 | Assembling Cooled LED Illuminator | Electromechanical assembly instructions for illuminator | osf.io/hywec [31] |
| Protocol 6 | Cutting and Drilling Clear Acrylic Sheet | Transparent orbital shaker platform build instructions | osf.io/69f8t [32] |
| Protocol 7 | Assembling Algal Shaker | Final assembly and testing steps | osf.io/ewc87 [33] |
| Protocol 8 | Measuring PPFD on Algal Shaker | Light output calibration protocol | osf.io/va54p [34] |

The above protocols explain in detail the steps required to build and test the illuminated orbital shaker.

## 6. Operation instructions

The illuminated orbital shaker offers three variable parameters: (1) platform shaking frequency, (2) growth light intensity, and (3) daily ratio of light and dark periods. These parameters can be operated independently. Their meaning and use is explained in the rest of this section.

The platform shaking frequency is set on the orbital shaker as described in its instruction manual. Usually, this would be done by turning a knob with a scale calibrated in revolutions per minute (RPM). The orbital shaker described in this manuscript supports frequencies between 0 and 210 RPM. The shaking speed needs to be high enough to keep the cell culture suspended and well mixed. Frequency higher than required unnecessarily increases the power consumption, heat generation, noise, and the risk of fall of flasks from the shaking platform. The validation experiments in this paper had the orbital shaker operating at 100 RPM.

The growth light intensity is controlled through the LED controller. There are two modes of operation: the trickle current and variable current modes, selected by a toggle switch. In the trickle current mode, a current of 26 mA is flowing through the LEDs, which creates photosynthetic photon flux density (PPFD) of ≈ 26 μmol · m^−2^ · s^−1^ inside a glass Erlenmeyer flask containing deionized water (Figure 3). The trickle current setting is used for maintaining slow growing cultures of cells or for microalgae species requiring low light conditions. The variable current mode allows adjusting the LED current between 240 mA and 1 A and PPFD between 220 and 800 μmol · m^−2^ · s^−1^. In the variable current mode, the LED current, and thus PPFD, are adjusted in seven steps by the rotary switch on the LED controller (Figure 3). The LED current is set by the resistors in the LED controller circuit and will remain the same irrespective of the implementation of the illuminated orbital shaker. The PPFD will depend on the geometry of the orbital shaker platform, the material of the anti-slip mat holding the culture flasks, spacing of the LED strips, and the type of the LED strips. It can therefore vary significantly in different implementations of the illuminated orbital shaker and will have to be calibrated individually.

**Figure 3:**
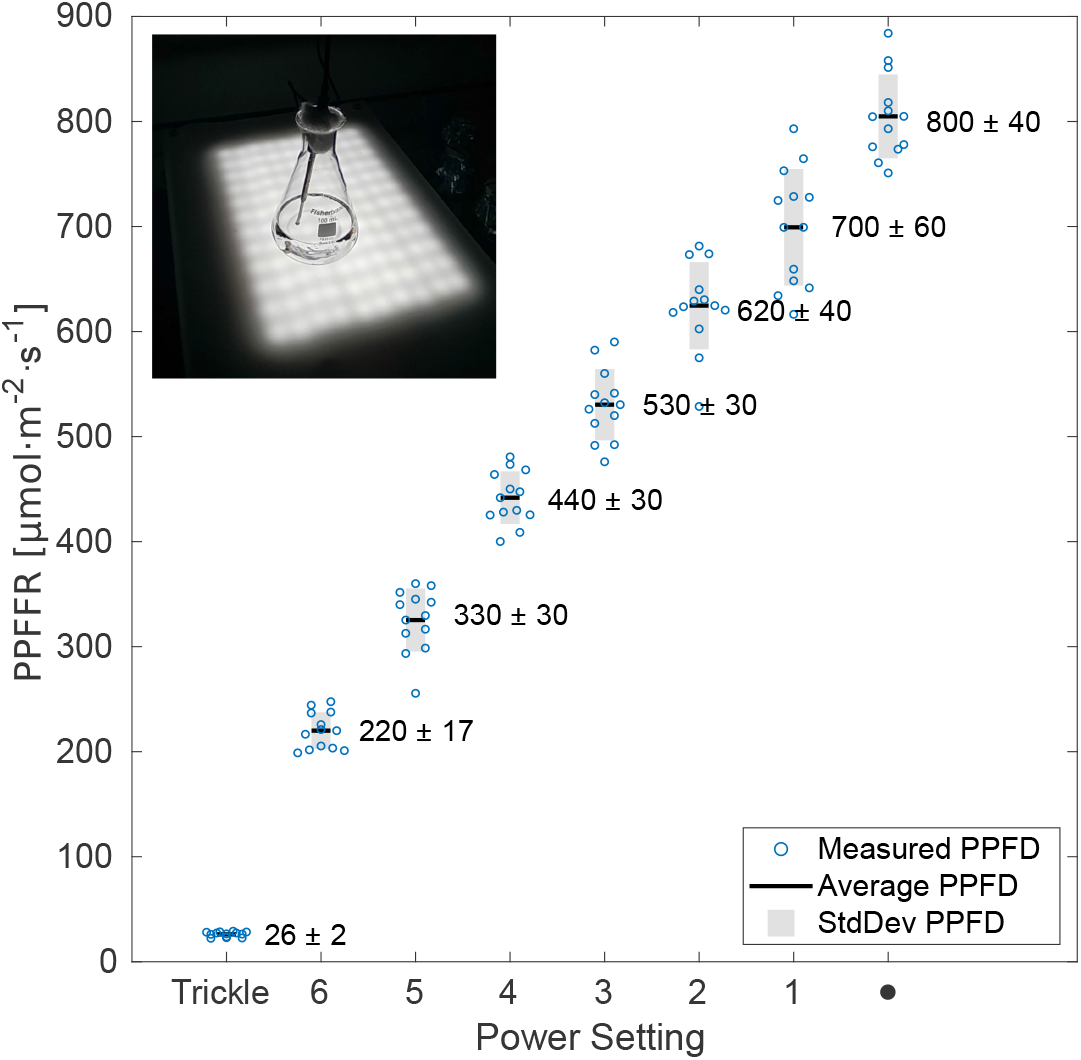
Calibration graph of photosynthetic photon flux density (PPFD) in water in relation to eight different LED current settings. For each setting, PPFD measurement was repeated in twelve different randomly selected positions on the illuminated platform. (inset) The light sensor was fixed vertically in deionized water inside 100 ml glass Erlenmeyer flask, to mimic the light conditions experienced by the microalgae culture. The graph shows the twelve measurements 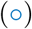 for each LED current setting, their average values (─), and standard deviations (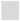, StdDev).

The growth light/dark cycle is adjusted by a 24-hour socket timer. By setting the timer to a night period, when the light is off, and a day period, when the light is on, the growth light can be programmed to mimic natural growing conditions. When this diurnal illumination is used, the cells in the culture become synchronized over time, assimilating, growing, and dividing in similar time periods of the day. This synchronization is beneficial to some experiments. When synchronization is not required, the light can be left turned on constantly for faster growth. In that situation, the culture growth is constant, with cells passing through the cell cycle unsynchronized.

The operation of the illuminated orbital shaker requires several safety considerations. The main risks are associated with the mains electricity, fire, and bright light. Ensure that the mains electrical leads are away from the orbital shaker to prevent the orbital platform from cutting through the insulation material of the cables. Make sure that mains socket connections are away from the orbital shaker. This will minimize the risk of fire or electrocution in case any flask containing the liquid culture falls from the orbital and spills its content. Ensure the orbital shaker is placed in a tidy space, away from splashing water, allow enough space for sufficient airflow to prevent overheating and increased risk of fire. Regularly check that the electrical cables and connections are sound and reliable. Check that the heatsink is not becoming clogged with dust, impeding good airflow. Fix unreliable or failing wiring and brush any dust off the heatsink whenever problems are identified. The orbital shaker must not move during operation even at the highest speed setting. In case of any movement, change its placement or place a sticky mat underneath to minimize the risk of falling. The illuminated orbital shaker is a source of bright light. Avoid staring at the illuminator. Use protective glasses if eye irritation occurs or if it is required by local regulations. Further safety instruction are listed in Supplementary Section S2.

## 7. Validation and characterization

The illuminated orbital shaker has been characterized and validated by measuring produced photosynthetic photon flux density (PPFD) and successful growth of microalgae cell cultures. PPFD has been measured using a calibrated light detector at randomly selected positions on the surface of the illuminated shaking platform and submerged in water inside glass Erlenmeyer flasks. Cell counting was done regularly to assess the increase in the density of the culture. The validation by microalgae growth was performed with two commonly used species of freshwater microalgae, *D. quadricauda* and *C. vulgaris*. The microalgae cultures were grown in different media at two different LED current settings and regularly evaluated by counting the cell density. The average cell doubling time was estimated for each species and growth condition. The experiments verified that the illuminated orbital shaker can be used for reliable cultivation of microalgae (Section 7.2 and Supplementary Section S1).

### 7.1 Photosynthetic photon flux density

This section discusses the calibration measurements of the light source in the illuminated orbital shaker. The light source is an array of white LEDs. The spectral density of a white LED is a compound of blue LED emission and green-red phosphorescence [19]. The white LED spectrum spans most of the photosynthetically active radiation (PAR) spectral range (400 nm − 700 nm) [35], meaning it can efficiently drive photosynthesis. The photosynthetic quantum yield in microalgae is a complex function of the wavelength. The combined complexity of the LED emission spectrum and the photosynthetic quantum yield spectral dependence requires specialized techniques for quantifying light sources for their ability to drive photosynthesis. Photosynthetic photon flux density (PPFD) is a measure of the number of photons, with the wavelength between 400 nm and 700 nm, crossing a unit area per unit of time [35]. PPFD is measured using a quantum sensor, which is a spectrally corrected light sensor with constant spectral sensitivity in the PAR region and a zero response outside. PPFD is typically expressed in μmol (photons) · m^−2^ · s^−1^.

A spherical micro quantum sensor (US-SQS, Waltz) was used in conjunction with a light meter (LI-250A, Li-COR). PPFD was measured in air, on the surface of the illuminated orbital shaker platform, and submerged in deionized water inside a glass Erlenmeyer flask, normally used for microalgae cultivation. The LED controller setting was switched between all eight supported LED current settings. The measurement was taken at twelve different randomly chosen positions to sample the PPFD across the illuminated orbital shaker platform area. The average value and the standard deviation were calculated for each medium and LED current setting (Table S2). The dependence of PPFD on the LED controller setting, measured in water, is plotted in Figure 3 and, measured in air, is in Supplementary Figure S2. The measurements show the expected increase in PPFD with the growing LED current as the power settings were changed. For each power setting, the twelve measurements are spread around their average value. The spread is due to the spatial inhomogeneity of the illumination, with local maxima right above the LEDs and minima in between the LEDs. This variation should have negligible effect on the microalgae cultures. The cell suspensions are being continuously mixed and therefore individual cells experience the same average PPFD over time. PPFD was slightly lower in water than air, which is consistent with the absorption and reflection losses introduced by the extra layers of glass and water (Supplementary Table S2). The PPFD can be set between 26 and 800 μmol · m^−2^ · s^−1^ (in water), which covers a broad range of radiant fluxes comparable to a wide range of natural daylight conditions. The protocol, detailing the experimental execution of the PPFD measurements and the data analysis steps, is at osf.io/va54p [34]. The raw measurement data, Matlab code for their analysis, and the resulting figures are available at osf.io/zamnf and in a GitHub repository at rebrand.ly/fmkm7hv.

### 7.2 Microalgae Cultivation

This section discusses microalgae cultivation using the bottom-illuminated orbital shaker. The objectives were to demonstrate the successful and consistent growth of microalgae at different PPFDs, to test microalgae growth in different media, and to find optimal cultivation conditions for future experiments. Two commonly used freshwater species, *Chlorella vulgaris* (#256, CCALA, Třeboň, Czech Republic) and *Desmodesmus quadricauda* (#463, CCALA), were cultivated. The experiments are described in further detail in Supplementary Section S1, introducing the experimental steps, protocols (Supplementary Table S1), and data analysis. This section reports on the results of an experiment, in which microalgae were grown at two different LED illuminator settings (PPFD of 26 and 220 μmol · m^−2^ · s^−1^) and in different growth media. Self-prepared medium ½SŠ [36] and a commercial Bold’s basal medium [37, 38] (BBM) (B5282, Merck, Gillingham, UK) were used. The ½SŠ medium was supplemented with 0.83 mM NaHCO_3_, as an additional source of carbon. The BBM medium was used both with 10 mM NaHCO_3_ and without any NaHCO_3_. Two light settings of 26 and 220 μmol · m^−2^ · s^−1^ were used. Cultures were seeded into fresh medium at the starting density of 2 × 10^5^ ml^−1^ for *D. quadricauda* and 1 × 10^6^ ml^−1^ for *C. vulgaris*. The cell densities were regularly obtained by counting using a Neubauer hemocytometer until saturation was reached (Supplementary Section S1.2).

#### 7.2.1 Cell Growth Rate Results

The cell density time evolution data for *D. quadricauda* are in Figure 4A and of *C. vulgaris* in Figure 4B. The graphs show the repeats of cell density counts for each day and experimental condition in the form of univariate scatter clouds (×, ∘, ∇, Δ). The average values of the cell densities are plotted as horizontal lines (─). The linear regression of the quasi-linear part of the plot for each condition are marked by the dotted lines (· · ·). The quasi-linear parts of the growth data were automatically identified and their linear regressions were calculated by a Matlab script (osf.io/52348 and on GitHub at rebrand.ly/t4zwgz1). The resulting doubling times during the exponential growth phase for each species and cultivation condition are listed in Table 3. The data show that doubling times shorten, and thus the growth rates become faster, for both species with brighter light (higher PPFD). The doubling times for *C. vulgaris* were similar in all three media (½SŠ, BBM, and BBM with NaHCO_3_). *D. quadricauda* grown in the ½SŠ medium had shorter doubling time and faster growth compared to when grown in the BBM medium^2^.

**Figure 4:**
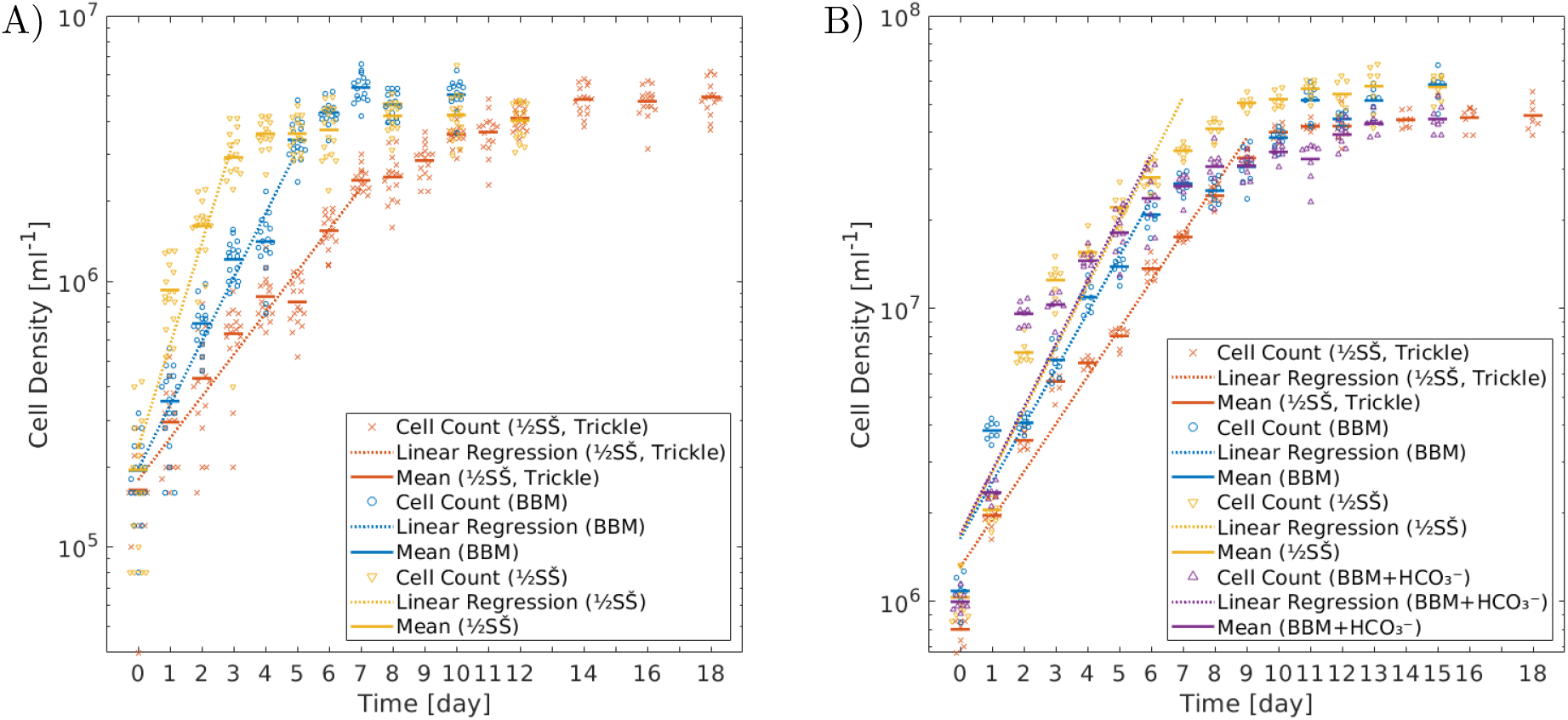
Growth data of (A) *D. quadricauda* and (B) *C. vulgaris* on semi-logarithmic plots. Cells were grown in (orange) ½SŠ medium at the trickle current setting (26 μmol · m^−2^ · s^−1^), (blue) ½SŠ at switch position 6 (220 μmol · m^−2^ · s^−1^), (yellow) BBM at switch position 6, and (purple) BBM with NaHCO_3_ at switch position 6. The average values of the cell density for each day and condition are marked by the colored horizontal bars (─). The cell density counted in each segment of the hemocytometer is plotted as a univariate scatter plot surrounding the average value (×, ∘, ∇, Δ). The linear regression of the quasi-linear part of the growth curve is marked by the colored dotted lines (…).

**Table 3:** List of doubling periods for *D. quadricauda* and *C. vulgaris* grown at PPFD of 26 μmol · m^−2^ · s^−1^ and 220 μmol · m^−2^ · s^−1^ in ½SŠ with 0.83 mM NaHCO_3_, BBM, and BBM with 10 mM NaHCO_3_. The listed doubling time values are the slopes of the linear regressions through the quasi-linear part of the logarithm of the cell culture density growth data. The stated uncertainties are the errors of the linear regression slope.

| Species | PPFD<br>[$\mu\text{mol} \cdot \text{m}^{-2} \cdot \text{s}^{-1}$] | Growth Medium | Doubling Time<br>[hours] |
| --- | --- | --- | --- |
| *D. quadricauda* | 26 (trickle current) | $\frac{1}{2}$SŠ + 0.83 mM $\text{NaHCO}_3$ | $46 \pm 3$ |
| *D. quadricauda* | 220 (switch position 6) | $\frac{1}{2}$SŠ + 0.83 mM $\text{NaHCO}_3$ | $19 \pm 3$ |
| *D. quadricauda* | 220 | BBM | $30 \pm 2$ |
| *C. vulgaris* | 26 | $\frac{1}{2}$SŠ + 0.83 mM $\text{NaHCO}_3$ | $44 \pm 2$ |
| *C. vulgaris* | 220 | $\frac{1}{2}$SŠ + 0.83 mM $\text{NaHCO}_3$ | $34 \pm 2$ |
| *C. vulgaris* | 220 | BBM | $37 \pm 2$ |
| *C. vulgaris* | 220 | BBM + 10 mM $\text{NaHCO}_3$ | $33 \pm 3$ |

The increase of the cell density over time did not follow perfect exponential model (linear increase on the semi-logarithmic plot) in these cultures. This had two reasons. Towards the saturation of the culture density, the growth naturally slowed down due to less light reaching the optically dense culture and the gradual exhaustion of nutrients. The second phenomenon was the daily fluctuation away from the ideal exponential growth model. We speculate, that the fluctuations were caused by a combination of hemocytometer loading errors and cell cycle oscillations in the synchronized cultures. After a period of stress^3^, the cultures did not exhibit steady exponential growth. Instead, the culture density increased considerably the first day after inoculation into fresh growth medium. This was followed by a day of subdued growth. These oscillations of faster and slower growth gradually evened out over the course of a week under stress-free conditions. Despite the cultures being conditioned for a week before the start of the experiments, the dilution to the initial inoculation density may have contributed to the observed daily fluctuations away from the ideal exponential growth model.

## 8. Capabilities of the Illuminated Orbital Shaker

- Growth area: 20 cm × 15 cm
- 100 ml Erlenmeyer flask capacity: 6
- T75 plastic tissue culture flask capacity: 2
- T25 plastic tissue culture flask capacity: 8
- Light source PPFD: 26 − 800 μmol · m^−2^ · s^−1^ (in eight steps, see Table S2)
- Light source peak power consumption: 29 W (excluding the shaker)
- Shaking speed: 0 − 210 RPM

## 9. Conclusions and Discussion

Protocols for building, testing, and using the bottom-illuminated orbital shaker for microalgae culture are presented. The protocols are accompanied by design files, which are editable using free-of-charge software. To encourage third-party customization for different purposes or experimental requirements, the design files, protocols, and code are released under minimally restrictive licenses^4^.

The illuminated orbital shaker light output was calibrated by measuring the photosynthetic photon flux density (PPFD). Its function for the intended purpose was verified by culturing *C. vulgaris* and *D. quadricauda*. The manuscript is accompanied by a set of protocols detailing the process of microalgae cultivation (Supplementary Section S1 and Table S1). The protocols introduce the growth media, the cultivation process, and the assessment of culture growth. The raw data from counting cell density and the Matlab code for their analysis are also openly shared.

The bottom-illuminated orbital shaker for microalgae culture was found perfectly suitable for our research on photosynthesis. Any suitably equipped workshop should be able to reproduce it within days, once all components are purchased, at a cost of around £300. All adopters are encouraged to openly share their applications, implementations, and changes to the illuminated orbital shaker. The remaining discussion covers possible improvements and modifications.

The design is flexible and well suited to alteration. A different orbital shaker can be used instead of the one described in this manuscript. The illuminated growth area and the maximum PPFD can be modified for specific needs. This requires scaling to a different-sized heatsink and/or changing the spacing and length of the LED strips. The light source color could be varied by choosing triple-color LED strips with three copies of the LED controller regulating each color separately. Longer than 24-hour light cycles can be achieved by using a weekly, rather than daily programmable timer. Alternatively, flexible control of timing and brightness can be done by driving the LED controller input from a microcontroller, as described below.

To support advanced and programmable growth protocols, the LED controller features an external control input. This allows the control of the growth light brightness by a microcontroller (e.g. Arduino or Raspberry Pi). The external input accepts either an analog voltage (0.30 V − 1.25 V), or a pulse-width modulation (PWM) with the amplitude of 1.25 V. The rotary switch can be conveniently used to match the maximum input voltage of the LED controller (1.25 V) to different microcontroller output voltages^5^ (e.g. 3.3 V or 5 V).

The illuminated orbital shaker could be placed inside a growth cabinet to allow temperature, humidity, and/or CO_2_ control for experiments or species requiring particular environmental conditions. ‘OpenTCC’ is an OSHW temperature-controlled cabinet that could be built to house the illuminated orbital shaker and maintain the flasks with microalgae cultures at a set temperature [39]. ‘Polar Bear’, a temperature- and humidity-controlled environmental chamber, is described in [40]. For microalgae cultivation, the humidity control could be left out of the ‘Polar Bear’ for simplicity. A well described guide to repurposing a refrigerator into a temperature-controlled environmental chamber is described on the commercial ‘BrewPi’ project website [41]. The design instructions and source code for the ‘BrewPi’ are open source. The company offers components for sale, which could simplify and speed up its the customization for microalgae cultivation. A large scale temperature-controlled walk-in space made from inexpensive parts is described in [42]. This could accommodate many illuminated orbital shakers for mass batch cultivation.

There are a number of OSHW solutions for CO_2_ control. A temperature- and CO_2_-controlled tissue culture cabinet, described by Pelling et al. [43], could be repurposed to house the illuminated orbital shaker. The described system lacks cooling capability, which limits its application without further redesign. A related company was started with the promise to sell a refined low-cost open source incubator based on the design by Pelling at al. It currently provides open source Arduino code for CO_2_ and temperature control on GitHub [44]. Another implementation of a temperature- and CO_2_-controlled incubator with design files and source code is described in a thesis by Al-Sayagh [45]. This device is too small to house the illuminated orbital shaker, but its low-cost closed-loop CO_2_ regulator can be potentially adapted for use with a larger cabinet and gas cylinder. A small temperature- and CO_2_-controlled incubator is described as a part of a live-cell imaging system in [46]. The manuscript discusses the regulation in detail, however the design files and source code are not open source and are available upon request only. A different implementation uses a decommissioned neonatal infant incubator [47]. Finally, a temperature- and CO_2_-controlled cabinet, using chemically generated CO_2_ rather than compressed gas, is described in an entry to the IGEM 2019 competition [48]. To the best of our knowledge, there is no existing OSHW cooled and CO_2_-controlled cabinet, which would be ideal for housing the illuminated orbital shaker.

The calibration between the LED controller setting and the PPFD at the surface of the LED illuminator was measured using a commercial PAR meter. This, or a similar device, may not be universally available, requiring an alternative solution for its accurate calibration. PARduino is an OSHW data logger combining a commercial quantum PAR sensor with an Arduino [49]. Kuhlgert et al. published an advanced portable OSHW plant phenotyping station, which incorporates a PAR sensor [50]. They used a low-cost 4-channel spectrally-resolved photodiode array instead of a quantum sensor and derived a transformation to convert its output voltages into PPFD. They verified the accuracy of their PAR sensor under different experimental conditions using a calibrated quantum sensor. Kutschera and Lamb simplified their design to build a stand-alone PAR meter based on an Arduino and the same photodiode array [51]. There are many other non-scientific PAR detector projects created by aquarium enthusiasts and hydroponic growers available online. Among them, one suggests using a particular low-cost lux meter as an approximate PAR meter after this was calibrated against a proprietary quantum sensor with different types of light sources [52]. PAR meters that do not rely on calibrated quantum sensors may offer a suitable low-cost alternative to proprietary solutions. However, their calibration, reproducibility, and accuracy need to be evaluated, given the specific experimental requirements and lighting conditions.

## 11. Human and animal rights

Not applicable

## 12. Acknowledgements

We would like to acknowledge the funding from Biotechnology and Biological Sciences Research Council (UK), Grant/Award Number: BB/R004803/1. We would like to thank the following people. Ladislav Nedbal of Forschungszentrum Jülich, Germany advised us on setting up the cultures and critically read the manuscript. Ben Blackburn of King’s College London operated the 3D printer to produce the LED controller case. Giorgio Perin and Patrick Jones of Imperial College London, United Kingdom lent us their quantum light sensor and light meter for the PPFD calibration measurements of the illuminated orbital shaker.

## Footnotes

1 Any changes to the total length of the LED strip must be accompanied by verifying that the maximum current delivered by the LED controller does not exceed the maximum current rating for the given length of the LED strip. Replace LED driver DC1 (Supplementary Figure S1) with LDU2430S700, LDU2430S600 or LDU2430S500 for maximum LED current of 700 mA, 600 mA or 500 mA, respectively. With longer LED strips, offering higher maximum current rating, wire them into two or more parallel sections, each powered by its own LED controller.

2 *D. quadricauda* was not cultured in BBM with NaHCO_3_ during these experiments, as large proportion of coenobia were malformed or had non-canonical cell number, when grown in this medium (data not shown).

3 Growth to saturation, large step change in the illumination light PPFD, more than 10-fold dilution with fresh medium.

4 The restrictions on the CC BY and BSD licenses require crediting the authors in any future use. The CC BY-SA license additionally mandates sharing any derivatives under a similar license.

5 Switch position 5 allows connecting the LED controller to a 3.3 Volt microcontroller and position 6 to a 5 Volt microcontroller.

## References

[1] J. Masojídek, G. Torzillo, and M. Koblížek. Photosynthesis in Microalgae. In: A. Richmond and Q. Hu (Eds.), Handbook of Microalgal Culture. John Wiley & Sons, 2013. Chap. 2, 21–36. doi: 10.1002/9781118567166.ch2.

[2] A. Guedes and F. Malcata. Bioreactors for Microalgae: A Review of Designs, Features and Applications. In: P. G. Antolli and Z. Liu (Eds.), Bioreactors: Design, Properties and Applications. Nova Science Publishers, 2012, 1–52.

[3] O. Pulz. Photobioreactors: Production Systems for Phototrophic Microorganisms, Appl Microbiol Biotechnol 57 (3 2001), 287–293. doi: 10.1007/s002530100702.

[4] H. S. Kim, T. L. Weiss, H. R. Thapa, T. P. Devarenne, and A. Han. A Microfluidic Photobioreactor Array Demonstrating High-Throughput Screening for Microalgal Oil Production, Lab Chip 14 (8 2014), 1415–1425. doi: 10.1039/C3LC51396C.

[5] C. Westerwalbesloh, C. Brehl, S. Weber, C. Probst, J. Widzgowski, A. Grünberger, C. Pfaff, L. Nedbal, and D. Kohlheyer. A Microfluidic Photobioreactor for Simultaneous Observation and Cultivation of Single Microalgal Cells or Cell Aggregates, PLoS One 14(4) (2019), 1–13. doi: 10.1371/journal.pone.0216093.

[6] M. A. Borowitzka. Commercial Production of Microalgae: Ponds, Tanks, Tubes and Fermenters, J Biotechnol 70(1) (1999). Biotechnological Aspects of Marine Sponges, 313–321. doi: 10.1016/S0168-1656(99)00083-8.

[7] A. P. Carvalho, L. A. Meireles, and F. X. Malcata. Microalgal Reactors: A Review of Enclosed System Designs and Performances, Biotechnol Prog 22(6) (2006), 1490–1506. doi: 10.1021/bp060065r.

[8] Q. Wang, H. Peng, and B. T. Higgins. Cultivation of Green Microalgae in Bubble Column Photobioreactors and an Assay for Neutral Lipids, JOVE 143 (2019), e59106. doi: 10.3791/59106.

[9] L. Rodolfi, G. Chini Zittelli, N. Bassi, G. Padovani, N. Biondi, G. Bonini, and M. R. Tredici. Microalgae for Oil: Strain Selection, Induction of Lipid Synthesis and Outdoor Mass Cultivation in a Low-Cost Photobioreactor, Biotechnol Bioeng 102(1) (2009), 100–112. doi: 10.1002/bit.22033.

[10] O. Savchenko, J. Xing, X. Yang, Q. Gu, M. Shaheen, M. Huang, X. Yu, R. Burrell, P. Patra, and J. Chen. Algal Cell Response to Pulsed Waved Stimulation and its Application to Increase Algal Lipid Production, Sci Rep 7 (2017), 42003. doi: 10.1038/srep42003.

[11] E. O. Ojo, H. Auta, F. Baganz, and G. J. Lye. Engineering Characterisation of a Shaken, Single-Use Photobioreactor for Early Stage Microalgae Cultivation using *Chlorella sorokiniana*, Bioresour Technol 173 (2014), 367–375. doi: 10.1016/j.biortech.2014.09.060.

[12] E. O. Ojo, H. Auta, F. Baganz, and G. J. Lye. Design and Parallelisation of a Miniature Photobioreactor Platform for Microalgal Culture Evaluation and Optimisation, Biochem Eng J 103 (2015), 93–102. doi: 10.1016/j.bej.2015.07.006.

[13] R. Lehmann. NinjaPBR - The Photobioreactor for Cyanobacteria and Microalgae. 2015. url: https://github.com/roblehmann/NinjaPBR (accessed 06/26/2020).

[14] J. Sanabria et al. Open Source Bioreactor. 2020. url: https://github.com/hackuarium/bioreactor/ (accessed 06/26/2020).

[15] A. Wishkerman and E. Wishkerman. Application Note: A Novel Low-Cost Open-Source LED System for Microalgae Cultivation, Comput Electron Agric 132 (2017), 56–62. doi: 10.1016/j.compag.2016.11.015.

[16] B. T. Nguyen and B. E. Rittmann. Low-Cost Optical Sensor to Automatically Monitor and Control Biomass Concentration in Microalgal Cultivation, Algal Res 32 (2018), 101–106. doi: 10.1016/j.algal.2018.03.013.

[17] B. T. Nguyen and B. E. Rittmann. A Simple Turbidity Monitor and Control System for Microalgae. 2018. url: https://www.instructables.com/id/A-Simple-Turbidity-Monitor-and-Control-System-for-/ (accessed 06/26/2020).

[18] J. Büchs. Introduction to Advantages and Problems of Shaken Cultures, Biochem Eng J 7(2) (2001). Special Issue: Shaking Bioreactors, 91–98. doi: 10.1016/S1369-703X(00)00106-6.

[19] J. Cho, J. H. Park, J. K. Kim, and E. F. Schubert. White Light-Emitting Diodes: History, Progress, and Future, Laser Photonics Rev 11(2) (2017), 1600147. doi: 10.1002/lpor.201600147.

[20] The Nobel Prize in Physics 2014. 2014. url: https://www.nobelprize.org/prizes/physics/2014/prize-announcement/ (accessed 04/17/2020).

[21] S. E. Loftus and Z. I. Johnson. Reused Cultivation Water Accumulates Dissolved Organic Carbon and Uniquely Influences Different Marine Microalgae, Front Bioeng Biotech 7 (2019), 101. doi: 10.3389/fbioe.2019.00101.

[22] E. J. Kim, S. Kim, H.-G. Choi, and S. J. Han. Co-Production of Biodiesel and Bioethanol using Psychrophilic Microalga *Chlamydomonas sp.* KNM0029C Isolated from Arctic Sea Ice, Biotechnol Biofuels 13(1) (2020), 20. doi: 10.1186/s13068-020-1660-z.

[23] M. Morita, Y. Watanabe, and H. Saiki. High Photosynthetic Productivity of Green Microalga *Chlorella sorokiniana*, Appl Biochem Biotechnol 87 (3 2000), 203–218. doi: 10.1385/ABAB:87:3:203.

[24] E. Ono and J. Cuello. Carbon Dioxide Mitigation Using Thermophilic Cyanobacteria, Biosyst Eng 96 (2007), 129–134. doi: 10.1016/j.biosystemseng.2006.09.010.

[25] P. Schulze, L. Barreira, H. Pereira, J. Perales, and J. Varela. Light Emitting Diodes (LEDs) Applied to Microalgal Production, Trends Biotechnol 32 (2014), 422–430. doi: 10.1016/j.tibtech.2014.06.001.

[26] N. Forget, C. Belzile, P. Rioux, and C. Nozais. Teaching the Microbial Growth Curve Concept using Microalgal Cultures and Flow Cytometry, J Biol Educ 44(4) (2010), 185–189. doi: 10.1080/00219266.2010.9656220.

[27] J. Nedbal. Illuminated Orbital Shaker for Microalgae Culture, protocols.io (2020). doi: 10.17504/protocols.io.bdubi6sn.

[28] J. Nedbal. Procuring Parts for Algal Shaker, protocols.io (2020). doi: 10.17504/protocols.io.bdtwi6pe.

[29] J. Nedbal. Assembling LED Controller Electronics, protocols.io (2020). doi: 10.17504/protocols.io.bdiai4ae.

[30] J. Nedbal. 3D Printing Ccase for LED Controller, protocols.io (2020). doi: 10.17504/protocols.io.bdici4aw.

[31] J. Nedbal. Assembling Cooled LED Illuminator, protocols.io (2020). doi: 10.17504/protocols.io.bcrniv5e.

[32] J. Nedbal. Cutting and Drilling Clear Acrylic Sheet, protocols.io (2020). doi: 10.17504/protocols.io.bcueiwte.

[33] J. Nedbal. Assembling Algal Shaker, protocols.io (2020). doi: 10.17504/protocols.io.bdcdi2s6.

[34] J. Nedbal. Measuring PPFD on Algal Shaker, protocols.io (2020). doi: 10.17504/protocols.io.bdyxi7xn.

[35] K. J. McCree. Photosynthetically Active Radiation. In: O. L. Lange, P. S. Nobel, C. B. Osmond, and H. Ziegler (Eds.), Physiological Plant Ecology I: Responses to the Physical Environment. Berlin, Heidelberg: Springer Berlin Heidelberg, 1981, 41–55. doi: 10.1007/978-3-642-68090-8_3.

[36] V. Zachleder and I. Šetlík. Effect of Irradiance on the Course of RNA Synthesis in the Cell Cycle of *Scenedesmus quadricauda*, Biol Plant 24(5) (1982), 341–353. doi: 10.1007/BF02909100.

[37] H. C. Bold. The Morphology of *Chlamydomonas chlamydogama*, sp. nov. Bulletin of the Torrey Botanical Club 76(2) (1949), 101–108. doi: 10.2307/2482218.

[38] H. W. Bischoff and H. C. Bold. Phycological Studies. IV: Some Soil Algae From Enchanted Rock and Related Algal Species. University of Texas, 1963, 93.

[39] C. Sánchez, P. Dessí, M. Duffy, and P. N. L. Lens. OpenTCC: An Open Source Low-Cost Temperature-Control Chamber, HardwareX 7 (2020), e00099. doi: 10.1016/j.ohx.2020.e00099.

[40] J. M. Pearce. Open-Source Lab: How to Build Your Own Hardware and Reduce Research Costs. Elsevier Science, 2013.

[41] E. Jacobs. BrewPi: A Modern Brewery Controller. 2020. url: https://www.brewpi.com/ (accessed 06/30/2020).

[42] E. Martinez and S. J. Agosta. Budget-Limited Thermal Biology: Design, Construction and Performance of a Large, Walk-In Style Temperature-Controlled Chamber, J Therm Biol 58 (2016), 29–34. doi: 10.1016/j.jtherbio.2016.03.009.

[43] A. Pelling. DIY CO_2_ Incubator Bioreactor for Mammalian Cell Culture. 2014. url: https://www.pellinglab.net/post/diy-diy-co2-incubator-bioreactor-for-mammalian-cell-culture (accessed 06/30/2020).

[44] Incuvers Inc. Model 1 Tri-Gas Incubator. 2019. url: https://github.com/Incuvers-Inc/Model-1 (accessed 06/30/2020).

[45] Y. H. Al-Sayagh. A Highly Adaptive and Cost Effective Second Generation Incubator (SGI) towards Educational, Research and Clinical Processes. MSc Thesis. University of Nebraska-Lincoln, Biological Systems Engineering, 2014.

[46] M. P. Walzik, V. Vollmar, T. Lachnit, H. Dietz, S. Haug, H. Bachmann, M. Fath, D. Aschenbrenner, S. Abolpour Mofrad, O. Friedrich, and D. F. Gilbert. A Portable Low-Cost Long-Term Live-Cell Imaging Platform for Biomedical Research and Education, Biosens Bio-electron 64 (2015), 639–649. doi: 10.1016/j.bios.2014.09.061.

[47] M. Brinn, S. F. Al-Sarawi, T. Lu, B. J. C. Freeman, J. Kumaratilake, and M. A. Henneberg. A Portable Live Cell Culture and Imaging System with Optional Umbilical Bioreactor Using a Modified Infant Incubator, Preprints (2017), 2017010137. doi: 10.20944/preprints201701.0137.v1.

[48] Team Amazonas Brazil. CO_2_ Incubator: IGEM 2019 Competition Entry. 2019. url: https://2019.igem.org/Team:Amazonas-Brazil/Hardware (accessed 06/30/2020).

[49] H. R. Barnard, M. C. Findley, and J. Csavina. PARduino: A Simple and Inexpensive Device for Logging Photosynthetically Active Radiation, Tree Physiol 34(6) (2014), 640–645. doi: 10.1093/treephys/tpu044.

[50] S. Kuhlgert, G. Austic, R. Zegarac, I. Osei-Bonsu, D. Hoh, M. I. Chilvers, M. G. Roth, K. Bi,D. TerAvest, P. Weebadde, and D. M. Kramer. MultispeQ Beta: a tool for large-scale plant phenotyping connected to the open PhotosynQ network, Royal Society Open Science 3(10) (2016), 160592. doi: 10.1098/rsos.160592.

[51] A. Kutschera and J. Lamb. Light Meter for Measuring Photosynthetically Active Radiation, Am J Plant Sci 9(12) (2018), 2420–2428. doi: 10.4236/ajps.2018.912175.

[52] S. Torpey. PAR Meter Hack $30. 2020. url: https://www.youtube.com/watch?v=YiVVAePNtXo (accessed 06/30/2020).

