## Supplementary Information for "Bottom-Illuminated Orbital Shaker for Microalgae Cultivation"

**Title:** Supplementary Materials to Bottom-Illuminated Orbital Shaker for Microalgae Cultivation

### S1. Supplementary Information: Growth of Microalgae

This supplementary section discusses the application of the bottom-illuminated orbital shaker to microalgae cultivation. Two species, *Chlorella vulgaris* (#256, CCALA, Třeboň, Czech Republic) and *Desmodesmus quadricauda* (#463, CCALA), were grown in different conditions from dilute suspensions up to saturation. Cell counting was done regularly to assess the increase in the cell density of the culture. The objectives were to demonstrate the successful and consistent growth of microalgae on the illuminated orbital shaker at different PPFDs, to test microalgae growth in different media, and to find optimal growth conditions for future experiments. Below, we describe the culture conditions, the cell density counting, the growth curves, and culture doubling periods. The associated protocols are listed in Supplementary Table S1.

#### S1.1 Culture Conditions: Light and Growth Medium

Different cultivation conditions were created by changing the LED controller setting and the growth media. Microalgae were grown with two different LED controller settings: The trickle current setting ($26\,\mu mol \cdot m^{-2} \cdot s^{-1}$) and the rotary switch position 6 setting ($220\,\mu mol \cdot m^{-2} \cdot s^{-1}$). Two types of growth media were tested: modified ½SŠ [1] and modified Bold's basal medium [2, 3] (BBM) (B5282, Merck, Gillingham, UK). ½SŠ was used because of the positive past experience with this medium, BBM was chosen as a widely available alternative. Modified ½SŠ medium was prepared similar to the recipe in [1]. The ½SŠ growth medium composition is in Supplementary Table S8 and its preparation is described in the protocol at osf.io/wv5ts [4]. The protocol differs from [1] by diluting the nutrients 12 times and by adding 0.83 mM $NaHCO_3$, supplementing the atmospheric $CO_2$ dissolved in the medium. The pH was adjusted to 7.5 with KOH. BBM was prepared by diluting the 50× stock solution in deionized water, adding the optional 10 mM $NaHCO_3$, adjusting pH, and filter sterilizing. The pH of BBM without carbon was adjusted to 6.6 using KOH. The pH of BBM with 10 mM $NaHCO_3$ was adjusted to 7.0 using $H_2SO_4$[1]. The cultures were grown in 100 ml Erlenmeyer flasks (15409103, Fisher Scientific, Loughborough, UK). The flasks were first cleaned, capped with cotton wool plug and aluminium foil lid, and autoclaved according to the protocol at osf.io/eynxp [5]. The microalgae cultures were inoculated into the Erlenmeyer flasks in 20 ml of sterile growth medium at the following cell densities: *C. vulgaris* at $1.0 \cdot 10^6\,ml^{-1}$ and *D. quadricauda* at $2.0 \cdot 10^5\,ml^{-1}$. The details are in the protocol at osf.io/pxq48 [6]. The cultures were cultivated shaking at 100 RPM at room temperature (23 – 25 °C) with a diurnal cycle of 14 hours of light, followed by 10 hours of darkness.

We found that in the ½SŠ medium, but not BBM or BBM with $NaHCO_3$, cell deposits were forming on the glass Erlenmeyer flasks near the surface of the medium with both species[2]. Especially with *C. vulgaris* in ½SŠ, the deposits were composed of strongly adhering clumps of cells attached to the flask walls. Therefore, the cell cultures were resuspended for daily cell density counting by trituration with a 20 ml syringe and a 21G needle, instead of vortexing on a shaker, as done normally.

[1] *D. quadricauda* was not cultivated in BBM with $NaHCO_3$ during these experiments, as large proportion of coenobia were malformed or had non-canonical cell number, when grown in this medium (data not shown).

[2] The cell accumulation on the flask walls was observed in plastic tissue culture flasks, too.

#### S1.2 Cell Density Counting

Cell counting was performed using a Neubauer hemocytometer (AC1000, Hawksley, Lancing, UK) on an inverted phase contrast microscope (CK 2, Olympus) with a 10× long-working distance objective (MVC-10X, Newport Spectra-Physics, Didcot, UK). This method is laborious, but simple to set up, robust, and less prone to artefacts, compared to the alternatives [7]. It copes well with cell clumping, varying cell size, and counting individual cells in multicellular coenobia of *D. quadricauda*. The cell density was counted once a day, until the interval was increased to every two days, as the culture densities neared their saturation. The protocol on cell density counting is at osf.io/u5paz [8]. In brief, Hemocytometer Sidekick phone app [9] was used to record the counts. In *C. vulgaris*, the counting was repeated in four segments of both counting chambers of the Neubauer hemocytometer. Due to the high cell density of *C. vulgaris* cultures, instead of counting the cells in the entire segment of the chamber, only a sub-section, typically containing 150 – 400 cells, was used. The numbers of cells in these sub-sections were scaled to the full area of the segments to calculate the total cell density in the cell suspension. In *D. quadricauda*, the counting was done in eight outer segments of both chambers. Counting was repeated three times, to get the numbers of coenobia with two, four, and eight cells, respectively. Malformed or hollow appearing coenobia were not counted. The results of the cell density counting were processed using a custom Matlab script to produce graphs and estimates of cell doubling times during the exponential growth phase. The raw cell density data, the analysis script, and the resulting graphs are available for download from GitHub at rebrand.ly/t4zwgz1.

### S2. Supplementary Information: Safety Instructions

The following list of safety precautions must only be used as a guidance to develop local safety rules for the operation of the bottom-illuminated orbital shaker.

#### S2.1 Instructions Related to Electrical Hazards

⚠ Regularly check the integrity of cables, plugs, and cases housing any electrical parts. Ensure they are intact and always protect them from damaging environment. Do not operate equipment with any signs of damage.

⚠ Let only trained and qualified people install or review the installation of the illuminated orbital shaker.

⚠ Ensure electrical sockets used for the illuminated orbital shaker are appropriately grounded. Consider connecting the illuminated orbital shaker to the mains electricity through a residual-current circuit breaker.

⚠ Do not install the illuminated orbital shaker in places where splashes of water or high humidity pose a risk. Do not install it outdoors, near sinks, steam generating devices, greenhouses, busy lab benches, and other locations with a risk of exposure to water or high humidity.

⚠ Do not install the illuminated orbital shaker in corrosive, volatile, dusty, explosive, combustible or flammable atmospheric conditions.

⚠ Install the illuminated orbital shaker only on sturdy non-slippery surfaces. Install it at waist height on a clutter-free worktop. A fall could result in electrical hazard, fire, spill of liquid or injury.

⚠ Keep all cables organized and tidy away from floor, edges of tables, sharp edges, and the orbital shaker platform. Use tape or cable clips to secure cables in safe locations to avoid trip hazard, damage to the cable insulation, and exposure to water or damaging chemicals.

⚠ Turn off the switch and unplug the illuminated orbital shaker from the socket before any maintenance, repair, repositioning or adjustment.

⚠ Do not touch the device with wet or sweaty hands. Do not insert any metallic objects into its voids.

⚠ Disconnect the device from electricity if anything unusual about it is discovered, such as becoming warm, hot or sounding unusual.

⚠ Disconnect the device from electricity if not used for a long period of time.

#### S2.2 Instructions Related to Operational Hazards

⚠ Develop specific instructions if your microalgae or growth media are hazardous. All instructions in this publication assume that the microalgae grown on the illuminated orbital shaker and the growth media are not hazardous to human health or the environment.

⚠ Prepare and handle growth media in a safe manner. Minimize volumes of chemicals handled, use appropriate personal protection equipment and precautions. Do not expose yourself, people around you or the environment to any chemicals.

⚠ Do not dispose of live microalgae into the chemical waste sink. Add bleach, Virkon, or other appropriate inhibiting agents for a sufficient amount of time before disposing of any microalgae cultures down the sink. Mix them sufficiently to ensure all microalgae, including those trapped on the walls of the flasks, get exposed to the agent. Work safely, wearing personal protection equipment when handling bleach or other cell-inhibiting agent. Handle the chemicals with care, do not breath the fumes or dust from powder-based substances.

⚠ Dispose of broken glass safely and clear up all spillages. Cleaning, autoclaving, handling and storing the Erlenmeyer flasks can result in their breaking and potentially spilling of their content. Broken glass is sharp and especially dangerous if contaminated. Ensure that spillage is safely contained and cleaned using appropriate equipment and the glass is disposed of in a safe manner. Wear protective gloves when handling shards of glass and make sure none is left behind, potentially injuring others in the future. Contaminated glass cannot be handled as standard glass waste. Make sure it is disposed of as a sharp contaminated biological/clinical waste.

⚠ Avoid spilling microalgae cultures onto the illuminated orbital shaker. Microalgae cultures are liquid. Combined with the shaking motion of the orbital shaker, the risk of a flask tipping over and spilling onto the electrical appliance is not insignificant. To minimize the risk, ensure the orbital shaker is on a stable, non-slippery support. Do not place the flasks near the edge of the platform. Keep the nonslippery mat on the surface of the shaking platform clean and free of dust, which could otherwise limit its adhesive properties. Keep the area around the illuminated orbital shaker tidy and free of items that could fall and tip over the flasks containing the liquid cultures. If spillage occurs, immediately disconnect the electrical power by pulling the plug from the socket before wiping the spillage and assessing the damage.

⚠ Dispose of all contaminated consumables in a safe manner, wipe down all potentially contaminated surfaces regularly with 70 % denatured ethanol. Handling chemicals and microalgae during the preparation of media, cell culture inoculation or cell counting contaminates various laboratory equipment, surfaces, and consumables like serological pipettes, pipette tips, cleaning tissue, cotton wool, aluminium foil, hemocytometer, and gloves. Use biological or clinical waste disposal routes for disposable items and wipe surfaces clean with 70 % denatured ethanol.

⚠ Avoid repetitive strain injury. Pipetting and logging cell counts on a tally counter are repetitive tasks that can result in an injury. Make sure you use ergonomic equipment and hand position. Take frequent breaks from repetitive activities.

⚠ Illuminated orbital shaker produces bright light. While it should not cause lasting damage to eyes, observe safety precautions. Do not come close with your eyes to the glowing surface, do not stare into the light and/or wear sunglasses or other protective eyewear.

### S3. Supplementary Figures and Tables

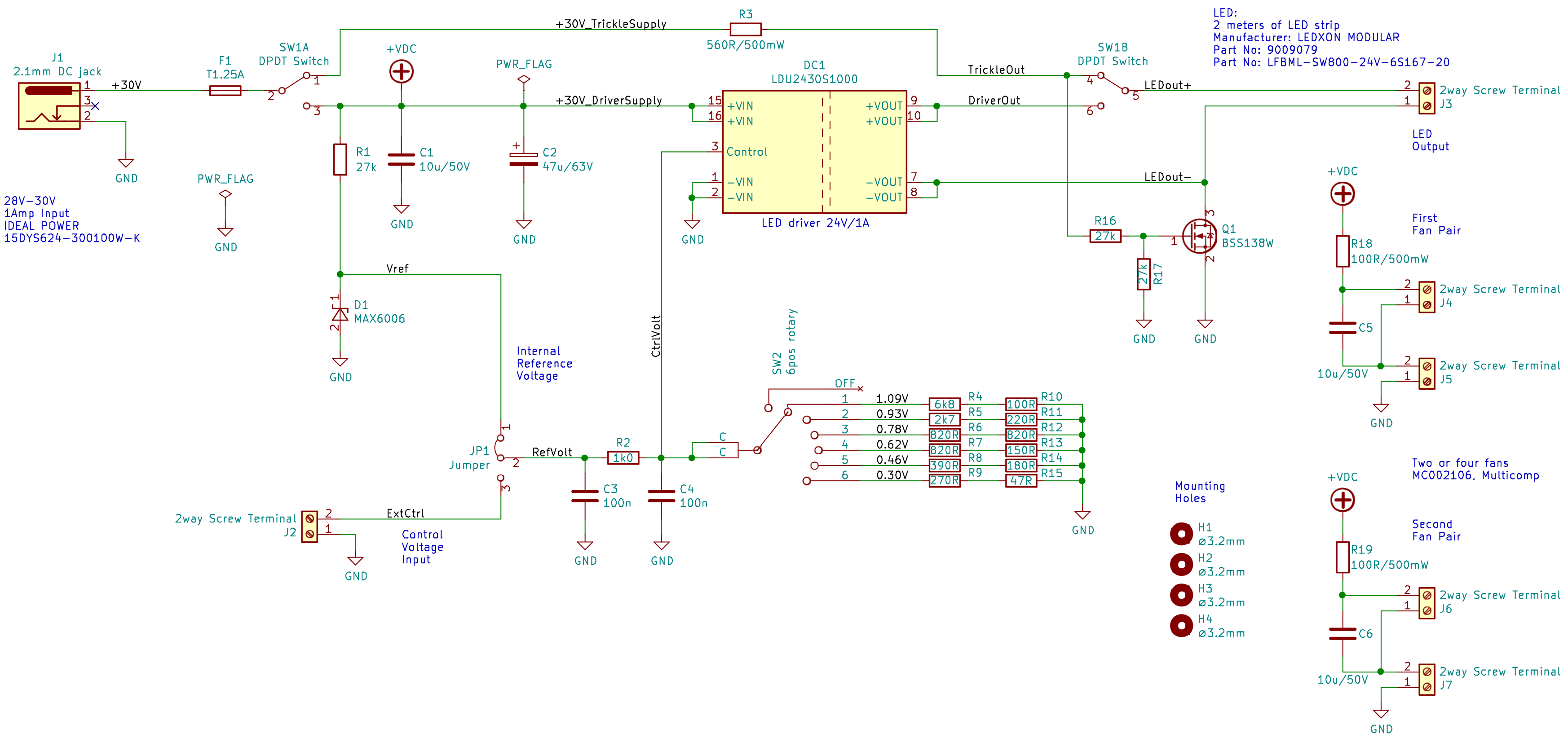


Figure S1: Electronic schematics of the LED controller. The LED controller uses a toggle switch (SW1) to alternate between trickle current, off-state and variable current modes. The variable current is controlled by rotary switch (SW2). Power (30 VDC/1 Amp) is supplied through a DC jack (J1). The LED illuminator is connected to a screw terminal (J3). The cooling fans are connected to screw terminals (J4 – J7). Optional external control input is through screw terminal (J2), when jumper (JP1) is in position 2-3. This input (J2) allows a microcontroller to regulate the LED brightness. More details about the electronics circuit are at osf.io/bftxm [10].

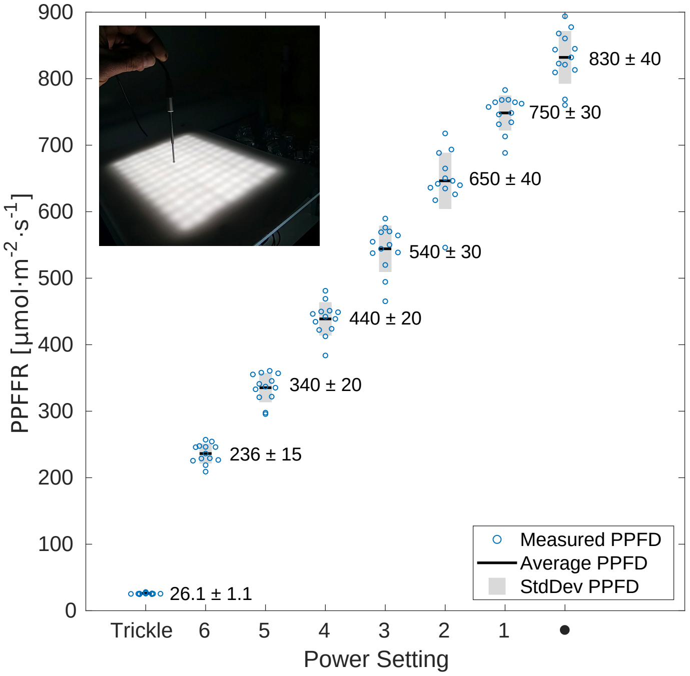


Figure S2: Calibration graph of photosynthetic photon flux density (PPFD) in air in relation to eight different LED current settings. For each setting, the measurement was repeated in twelve different randomly selected positions on the illuminated platform. (inset) The light sensor was held vertically in close contact with the illuminated platform. The graph shows the twelve measurements (○) for each LED current setting, their average values (—), and standard deviations (■, StdDev).

| Designator | Protocol | Summary | Protocol URL |
|---|---|---|---|
| Protocol S1 | Preparing ½SŠ Algal Inorganic Nutrient Medium | Growth medium recipe | osf.io/wv5ts [4] |
| Protocol S2 | Autoclaving Erlenmeyer Flasks for Sterile Microalgae Cultures | Cleaning and preparing flasks; autoclaving flasks | osf.io/eynxp [5] |
| Protocol S3 | Counting Microalgae Culture Density | Using hemocytometer to assess culture density | osf.io/u5paz [8] |
| Protocol S4 | Culturing *C. vulgaris* and *D. quadricauda* | Cell culture protocol | osf.io/pxq48 [6] |

Table S1: List of protocols with instructions on culturing microalgae. All protocols are released under CC BY 4.0 license. The URL links to the protocols in this table lead to their static snapshots, current at the publication time.

| **Switch Combination** | **LED Current** [**mA**] | **PPFD in Water** [$\mu$**mol** $\cdot$ **m**$^{-2}$ $\cdot$ **s**$^{-1}$] | **PPFD in Air** [$\mu$**mol** $\cdot$ **m**$^{-2}$ $\cdot$ **s**$^{-1}$] |
|---|---|---|---|
| Trickle | 26 | $26 \pm 2$ | $26.1 \pm 1.1$ |
| 6 | 241 | $220 \pm 17$ | $236 \pm 15$ |
| 5 | 363 | $330 \pm 30$ | $340 \pm 20$ |
| 4 | 492 | $440 \pm 30$ | $440 \pm 20$ |
| 3 | 621 | $530 \pm 30$ | $540 \pm 30$ |
| 2 | 745 | $620 \pm 40$ | $650 \pm 40$ |
| 1 | 873 | $700 \pm 60$ | $750 \pm 30$ |
| • | 1000 | $800 \pm 40$ | $830 \pm 40$ |

Table S2: Calibration data for the LED illuminator. The LED controller can be set to eight different light output settings. A toggle switch chooses between the Trickle current and seven different Variable current settings, selected by the rotary switch. The LED current, PPFD in water-filled flask, and PPFD in air on the surface of the shaking platform are listed for each switch combination. The PPFD is expressed as the average and the standard deviation of measurements in twelve different randomly selected positions on the illuminated shaker platform.

| Item | Qty | Description | Distributor | Manufacturer | Order Code | Part Number | Unit Price [£] | Total [£] |
|---|---|---|---|---|---|---|---|---|
| 1 | 1 | Orbital Shaker | eBay | | 283712826465 | KJ-201BD | 95.30 | 95.30 |
| **Total** | | | | | | | | **95.30** |

Table S3: Laboratory parts for the illuminated orbital shaker. The only laboratory part required to build the system is an orbital shaker. A low cost orbital shaker is being used here.

| Item | Qty | Value | Description | Distributor | Manufacturer | Order Code | Part Number | Unit Price [£] | Total [£] |
|---|---|---|---|---|---|---|---|---|---|
| 1 | 3 | 10u/50V | 10μF/50V X5R 1206 ceramic capacitor (pack of 5) | Farnell | Murata | 2672214 | GRT31CR61H106KE01L | 0.525 | 2.625 |
| 2 | 1 | 47u/63V | 47μF/63V electrolytic capacitor | Farnell | Panasonic | 2326166 | EEE1JA470UP | 0.679 | 0.679 |
| 3 | 2 | 100n | 100nF/50V X7R 0603 ceramic capacitor (pack of 10) | Farnell | Multicomp | 1759122 | MC0603B104K500CT | 0.0341 | 0.341 |
| 4 | 1 | MAX6006 | 1.25V shunt reference | Farnell | Maxim Integrated | 2511278 | MAX6006BEUR+T | 1.29 | 1.29 |
| 5 | 1 | LDU2430S1000 | 24V/1A adjustable LED driver | Farnell | XP Power | 1738296 | LDU2430S1000 | 8.23 | 8.23 |
| 6 | 1 | Fuse Holder | 5x20mm fuse holder (pack of 10) | Farnell | Schurter | 2309093 | 751.0052 | 0.103 | 1.03 |
| 7 | 1 | T1.25A | 1.25A slow-burning fuse (pack of 10) | Farnell | Eaton Bussmann | 1123242 | S506-1.25-R | 0.836 | 8.36 |
| 8 | 1 | 2.1mm DC Jack | 2.1mm 2A/16V DC power jack | Farnell | Clif Electronics | 2450496 | FC681478 | 1.84 | 1.84 |
| 9 | 6 | 2way Screw Terminal | 2way 3.81mm screw terminal | Farnell | Multicomp | 2007985 | MC000018 | 0.481 | 2.886 |
| 10 | 1 | 3way Shunt Header | 3way 2.54mm header (pack of 50) | Farnell | Multicomp | 1593412 | 2211S-03G | 0.0156 | 0.78 |
| 11 | 1 | Shunt Jumper | 2.54 mm shunt jumper (pack of 50) | Farnell | Multicomp | 2834673 | MC-2228CG | 0.0262 | 1.31 |
| 12 | 1 | BSS138W | 50V 200mA N-channel MOSFET (pack of 5) | Farnell | Diodes | 1713833 | BSS138W-7-F | 0.265 | 1.325 |
| 13 | 3 | 27k | 27kΩ 0603 1% SMD resistor (pack of 10) | Farnell | Multicomp | 2447315 | MCWR06X2702FTL | 0.0038 | 0.038 |
| 14 | 1 | 1k0 | 1kΩ 0603 1% SMD resistor (pack of 10) | Farnell | Multicomp | 2447272 | MCWR06X1001FTL | 0.0038 | 0.038 |
| 15 | 1 | 560R/500mW | 560Ω 1206 1% 500mW SMD resistor (pack of 10) | Farnell | TE Connectivity | 2332136 | CRGH1206F560R | 0.0447 | 0.447 |
| 16 | 1 | 6k8 | 6.8kΩ 0603 1% SMD resistor (pack of 10) | Farnell | Multicomp | 2447427 | MCWR06X6801FTL | 0.0034 | 0.034 |
| 17 | 1 | 2k7 | 2.7kΩ 0603 1% SMD resistor (pack of 10) | Farnell | Multicomp | 2447324 | MCWR06X2701FTL | 0.0034 | 0.034 |
| 18 | 3 | 820R | 820Ω 0603 1% SMD resistor (pack of 10) | Farnell | Multicomp | 2447437 | MCWR06X8200FTL | 0.0035 | 0.035 |
| 19 | 1 | 390R | 390Ω 0603 1% SMD resistor (pack of 10) | Farnell | Multicomp | 2447353 | MCWR06X3900FTL | 0.0037 | 0.037 |
| 20 | 1 | 270R | 270Ω 0603 1% SMD resistor (pack of 10) | Farnell | Multicomp | 2447314 | MCWR06X2700FTL | 0.0038 | 0.038 |
| 21 | 1 | 100R | 100Ω 0603 1% SMD resistor (pack of 10) | Farnell | Multicomp | 2447227 | MCWR06X1000FTL | 0.0034 | 0.034 |
| 22 | 1 | 220R | 220Ω 0603 1% SMD resistor (pack of 10) | Farnell | Multicomp | 2447298 | MCWR06X2200FTL | 0.0034 | 0.034 |
| 23 | 1 | 150R | 150Ω 0603 1% SMD resistor (pack of 10) | Farnell | Multicomp | 2447255 | MCWR06X1500FTL | 0.0035 | 0.035 |
| 24 | 1 | 180R | 180Ω 0603 1% SMD resistor (pack of 10) | Farnell | Multicomp | 2447267 | MCWR06X1800FTL | 0.0034 | 0.034 |
| 25 | 1 | 47R | 47Ω 0603 1% SMD resistor (pack of 10) | Farnell | Multicomp | 2694082 | MCWF06P47R0FTL | 0.0083 | 0.083 |
| 26 | 2 | 100R/500mW | 100Ω 1206 1% 500mW SMD resistor (pack of 10) | Farnell | TE Connectivity | 2332126 | CRGH1206F100R | 0.0423 | 0.423 |
| 27 | 1 | DPDT Switch | DPDT On-Of -On toggle switch | Farnell | Multicomp | 9473556 | 1MD4T1B1M1QE | 2.60 | 2.60 |
| 28 | 1 | 6pos Rotary Switch | 6-position rotary switch | Farnell | Nidec Copal | 2854809 | SS-10-16NP-LE | 2.48 | 2.48 |
| 29 | 1 | DIP-8 Socket | DIP-8 socket for rotary switch | Farnell | Multicomp | 2668408 | SPC15494 | 0.133 | 1.33 |
| **Total** | | | | | | | | | **38.45** |

Table S4: Electronics parts for the illuminated orbital shaker. Some of the parts are quite common and likely to be found in electronics workshops without the need to purchase them. Some of the electronics parts are subject to minimum order requirement and thus higher quantity has to be purchased than will be required.

| Item | Qty | Value | Description | Distributor | Manufacturer | Order Code | Part Number | Unit Price [£] | Total [£] |
|---|---|---|---|---|---|---|---|---|---|
| 1 | 4 | M4 Nut | M4 zinc-plated steel nut (pack of 100) | Farnell | TR Fastenings | 1419449 | M4-HFST-Z100- | | 1.59 |
| 2 | 8 | M4 Washer | M4 zinc-plated steel washer (pack of 100) | Farnell | TR Fastenings | 2506009 | DM4-FASTWAZ100DIN125 | | 1.14 |
| 3 | 4 | M4×12 Screw | M4 12mm zinc-plated pan head screw (pack of 100) | Farnell | TR Fastenings | 1419994 | M4 12 PRSTMC Z100 | | 2.08 |
| 4 | 4 | M3×10 Screw | M3 10mm countersunk zinc-plated screw (pack of 100) | Farnell | TR Fastenings | 1420398 | M3 10 KRSTMC Z100 | | 1.64 |
| 5 | 4 | 50mm M4 Standof | 50mm M4 standof male-female | Farnell | Ettinger | 1466738 | 05.14.501 | 0.73 | 2.92 |
| **Total** | | | | | | | | | **9.37** |

Table S5: Fixings for the illuminated orbital shaker. Widely available metric fixings are required and may be available in machine workshops. In world regions using imperial threads, the substitute parts are described in the protocol at osf.io/jy7gc [11].

| Item | Qty | Value | Description | Distributor | Manufacturer | Order Code | Part Number | Unit Price [£] | Total [£] |
|---|---|---|---|---|---|---|---|---|---|
| 1 | 2 | 1m LED Strip | LED strip, 1m, cool white, 24VDC, 14.4W | Farnell | Ledxon Modular | 2214009 | 9009079 | 25.84 | 51.68 |
| 2 | 4 | 40mm Fan | 12 VDC Fan, 60mA, 40mm | Farnell | Multicomp | 2816685 | MC002106 | 2.11 | 8.44 |
| 3 | 1 | 30V/1A AC Adaptor | 30V/1A 2.1mm DC power supply | Farnell | Ideal Power | 2771453 | 15DYS624-300100W-K | 23.37 | 23.37 |
| 4 | 1 | LED Heatsink | Heatsink, 200 mm x 150 mm x 40 mm, 0.5 °C/W | Farnell | Fischer Elektronik | 4621906 | SK 47/150 SA | 35.71 | 35.71 |
| 5 | 2 | Black Cable | 0.5 $mm^2$ black cable for LED strip connection | Farnell | Pro Power | 2528081 | PP001185 | 0.315 | 0.63 |
| 6 | 2 | Red Cable | 0.5 $mm^2$ red cable for LED strip connection | Farnell | Pro Power | 2528174 | PP001269 | 0.315 | 0.63 |
| 7 | 1 | Clear Acrylic | 4 mm clear acrylic sheet (any make, > 30cm×20cm) | RS | RS Pro | 824-660 | 824-660 | 32.72 | 32.72 |
| 8 | 1 | 24H Time Switch | Programmable 24-hour socket time switch (any make) | Farnell | Pro Elec | 2777066 | PEL00407 | 2.01 | 2.01 |
| **Total** | | | | | | | | | **155.19** |

Table S6: LED illuminator parts for the illuminated orbital shaker.

| Item | Qty | Value | Description | Distributor | Manufacturer | Order Code | Part Number | Unit Price [£] | Total [£] |
|---|---|---|---|---|---|---|---|---|---|
| **Workshop Tools** | | | | | | | | | |
| 1 | 1 | M3 Tap Set | M3 thread tap set (any make, #4-40 for imperial threads) | Farnell | Ruko | 375238 | 230-030 | 8.77 | 8.77 |
| 2 | 1 | Tap Wrench | Tap Wrench (any make) | Farnell | Ruko | 376395 | 241 001 | 10.71 | 10.71 |
| 3 | 1 | Tool Kit | Mechanical workshop tool kit (any make) | RS | RS Pro | 829-6561 | 829-6561 | 108.59 | 108.59 |
| 4 | 1 | DMM | General digital multimeter (any make) | RS | RS Pro | 123-1930 | 123-1930 | 30.00 | 30.00 |
| **Stationeries** | | | | | | | | | |
| 5 | 1 | Multipurpose Glue | All purpose clear adhesive (any make) | Amazon | Bostik | B0001OZI48 | All Purpose | 1.58 | 1.58 |
| 6 | 1 | Scissors | Common scissors (any make) | Amazon | Helix | B00XP1V0UU | Oxford 13cm | 1.39 | 1.39 |
| 7 | 1 | Pen | Fine tip permanent marker pen (not black, any make) | Amazon | Staedtler | B005DPPQAG | 733449 | 1.66 | 1.66 |
| 8 | 1 | Ruler | Ruler 30 cm (any make) | Amazon | Q Connect | B000NMBTUK | Ruler | 0.59 | 0.59 |
| **Soldering Equipment** | | | | | | | | | |
| 9 | 1 | Tweezers | Watchmakers tweezers (any make) | Farnell | Duratool | 3127692 | 1PK-125T-F | 3.22 | 3.22 |
| 10 | 1 | Solder Flux | Solder f ux (any make) | Farnell | Chip Quik | 1850220 | SMD291NL | 11.94 | 11.94 |
| 11 | 1 | Solder Wire | Thin solder wire (any make) | Farnell | Duratool | 3262209 | D03341 | 5.18 | 5.18 |
| 12 | 1 | Soldering Station | Soldering station suitable for SMD (any make) | Farnell | Metcal | 1560738 | PS-900 | 187.00 | 187.00 |
| 13 | 1 | Electrical Tape | PVC Electrical Insulation tape (any make) | Farnell | Pro Power | 152346 | PVC TAPE 1920B | 1.18 | 1.18 |
| 14 | 1 | IPA | Isopropyl alcohol for cleaning (any make) | Amazon | Hexeal | B079YVPZDF | IPA | 6.79 | 6.79 |
| 15 | 1 | Small Tub | Margarine tub or something similar | | | | | | |
| 16 | 1 | Brush | Old toothbrush or stif paintbrush | | | | | | |
| **3D Printing** | | | | | | | | | |
| 17 | 1 | 3D Printer | FDM 3D Printer with build material (any make, or outsource) | | Stratasys | uPrint SE Plus | uPrint SE Plus | | |
| **Cutting and Drilling Tools (Either tools or laser cutter, both not required)** | | | | | | | | | |
| 18 | 1 | Drill | General hand or pillar drill (any make) | Amazon | Skil | B00IINANZ8 | 6221AB | 44.99 | 44.99 |
| 19 | 1 | 4mm Drill Bit | HSS drill bit 4 mm (any make, ideally spur-point bit for plastic) | Farnell | Ruko | 378124 | 201 040 | 0.44 | 0.44 |
| 20 | 1 | Hacksaw | 300 mm hacksaw (any make) | Farnell | Duratool | 2103261 | D02166 | 7.32 | 7.32 |
| 21 | 1 | P150 Sandpaper | P80-P180 sandpaper (any make) | Amazon | 3M | B001PNBC0I | 20150 | 4.49 | 4.49 |
| 22 | 1 | Laser Cutter | Optional laser cutter for plastics (any make) | | | | | | |
| **Total** | | | | | | | | | **435.84** |

Table S7: Tools used in the build of the illuminated orbital shaker. Most of these tools should be widely available and therefore should not need to be purchased. The last section **Cutting and Drilling Tools** lists both hand tools and a laser cutter to cut out the shaker platform out of clear acrylic. Only one set of these tools is required - either the hand tools or the laser cutter.

| **Component** | **Quantity** | **Final Molarity** | **Note** |
|---|---|---|---|
| Deionized water ($H_2O$) | 500 ml | N/A | Autoclaved |
| 1.83 M $KNO_3$<br>Potassium nitrate stock | 833 µl | 3.0 mM | |
| 174 mM $KH_2PO_4$<br>Monopotassium phosphate stock | 417 µl | 145 µM | |
| 82.8 mM $MgSO_4 \cdot 7\,H_2O$<br>Magnesium sulphate heptahydrate stock | 417 µl | 69 µM | |
| 500 mM $NaHCO_3$<br>Sodium bicarbonate stock | 833 µl | 833 µM | Do not autoclave |
| 10.9 mM $C_{10}H_{12}N_2NaFeO_8$<br>EDTA iron(III) sodium salt stock | 417 µl | 9.1 µM | Do not autoclave |
| 79.3 mM $CaCl_2$<br>Calcium chloride stock | 417 µl | 66 µM | |
| Trace elements stock | 42 µl | See table (b) | Do not autoclave |
| 2 M KOH<br>Potassium hydroxide solution | 20 µl | 80 µM | Adjust pH to 7.5 |

(a) ½SŠ inorganic growth medium recipe

| **Component** | **Quantity [mg·l$^{-1}$]** | **Stock Solution Molarity [mM]** |
|---|---|---|
| Boric acid ($H_3BO_3$) | 833 | 13.4 |
| Manganese chloride tetrahydrate ($MnCl_2 \cdot 4\,H_2O$) | 3290 | 16.6 |
| Zinc sulfate heptahydrate ($ZnSO_4 \cdot 7\,H_2O$) | 2680 | 9.31 |
| Ammonium molybdate tetrahydrate ($(NH_4)_6Mo_7O_{24} \cdot 4\,H_2O$) | 171 | 0.138 |
| Ammonium metavanadate ($NH_4VO_3$) | 14 | 0.120 |
| Cobalt sulfate heptahydrate ($CoSO_4 \cdot 7\,H_2O$) | 617 | 2.19 |
| Copper sulfate pentahydrate ($CuSO_4 \cdot 5\,H_2O$) | 945 | 3.78 |

(b) Trace elements recipe for the ½SŠ inorganic growth medium

Table S8: Recipe for ½SŠ inorganic growth medium. The medium is prepared in deionized water from salts providing nutrients to the microalgae. The recipe is designed for growing microalgae suspensions up to the biomass dry weight of $0.5\,g \cdot l^{-1}$. (a) The table lists the constituents to prepare 500 ml ½SŠ inorganic growth medium from stock solutions of salts. The final step involves adjusting pH to 7.5 by adding KOH. All solutions are initially filter sterilized by 0.2 µm syringe filters and aseptically added to the autoclaved deionized water in the listed order. (b) Recipe for preparing a stock solution of trace elements added to the ½SŠ inorganic growth medium in (a).

### References

[1] V. Zachleder and I. Šetlík. Effect of Irradiance on the Course of RNA Synthesis in the Cell Cycle of *Scenedesmus quadricauda*, *Biol Plant* 24(5) (1982), 341–353. DOI: 10.1007/BF02909100.

[2] H. C. Bold. The Morphology of *Chlamydomonas chlamydogama*, sp. nov. *Bulletin of the Torrey Botanical Club* 76(2) (1949), 101–108. DOI: 10.2307/2482218.

[3] H. W. Bischoff and H. C. Bold. Phycological Studies. IV: Some Soil Algae From Enchanted Rock and Related Algal Species. University of Texas, 1963, 93.

[4] J. Nedbal and L. Gao. Preparing 1/2 SŠ Algal Inorganic Nutrient Medium, *protocols.io* (2020). DOI: 10.17504/protocols.io.bdzai72e.

[5] J. Nedbal. Autoclaving Erlenmeyer Flasks for Sterile Algal Cultures, *protocols.io* (2020). DOI: 10.17504/protocols.io.bd2di8a6.

[6] J. Nedbal. Culturing *Chlorella vulgaris* and *Desmodesmus quadricauda*, *protocols.io* (2020). DOI: 10.17504/protocols.io.bd6vi9e6.

[7] N. R. Moheimani, M. A. Borowitzka, A. Isdepsky, and S. F. Sing. Standard Methods for Measuring Growth of Algae and their Composition. In: *Algae for Biofuels and Energy*. Springer Netherlands, 2012, 265–284. DOI: 10.1007/978-94-007-5479-9_16.

[8] J. Nedbal. Counting Microalgae Culture Density, *protocols.io* (2020). DOI: 10.17504/protocols.io.bd7wi9pe.

[9] M. Fuentes. Hemocytometer Sidekick App. 2018. URL: https://www.hemocytometer.org/hemocytometer-sidekick-app/ (accessed 02/07/2020).

[10] J. Nedbal. Assembling LED Controller Electronics, *protocols.io* (2020). DOI: 10.17504/protocols.io.bdiai4ae.

[11] J. Nedbal. Procuring Parts for Algal Shaker, *protocols.io* (2020). DOI: 10.17504/protocols.io.bdtwi6pe.
